# Within-patient evolution to piperacillin/tazobactam resistance in a clinical isolate of *Escherichia coli* due to IS*26*-mediated amplification of *bla*_TEM-1B_

**DOI:** 10.1101/2020.02.28.969360

**Authors:** Alasdair T. M. Hubbard, Jenifer Mason, Paul Roberts, Christopher M. Parry, Caroline Corless, Jon van Aartsen, Alex Howard, Alice J. Fraser, Emily R. Adams, Adam P. Roberts, Thomas Edwards

**Author notes:** Corresponding authors: Dr. Alasdair Hubbard, Dr. Thomas Edwards.

## Abstract

A novel phenotype of *Escherichia coli* and Klebsiella pneumoniae resistant to piperacillin/tazobactam (TZP), but susceptible to carbapenems and 3^rd^ generation cephalosporins has recently emerged. The resistance mechanism of this phenotype has been identified as hyperproduction of the β-lactamase *bla*_TEM_, however the mechanism of hyperproduction in isolates lacking promoter region mutations is not well understood. We sought to understand this mechanism by focussing on a pair of isolates obtained from an individual patient across two infection episodes and displaying within-patient evolution to TZP resistance. Following confirmation that the two isolates were clonal, we found that the TZP-resistant isolate hyperproduced a β-lactamase but lacked mutations within β-lactamase promoter regions. Hybrid assembly of long and short sequencing reads of the two isolates revealed both harboured a novel IS*26*-flanked composite transposon containing several antibiotic resistance genes, including *bla*_TEM-1B_, which was designated Tn*6762*. These resistance genes are also found to be present on a translocatable unit which had excised from Tn*6762* in the TZP-resistant isolate. By replicating the evolutionary event leading to TZP resistance we were able to observe excision of the translocatable unit from Tn*6762* following exposure to TZP and capture the TU in a plasmid containing a copy of IS*26*. Subsequent amplification of the TU, and by extension *bla*_TEM-1B_, leads to β-lactamase hyperproduction and TZP resistance. Despite a significant increase in gene copy number (P value = <0.0001), we found that the TZP-resistant isolate was as fit as the susceptible ancestor. This mechanism of gene amplification, and the subsequent hyperproduction, of *bla*_TEM-1B_ is an important consideration when using genomic data to predict resistance/susceptibility to TZP.

## Introduction

β-lactam/β-lactamase inhibitor combinations were developed to overcome the activity of class A (e.g. TEM-1) and class C β-lactamases (BLs) (e.g. AmpC) [1, 2], which inactivate β-lactam antibiotics by hydrolysing the β-lactam ring. New β-lactamase inhibitors, particularly metallo-β-lactamase inhibitors, are still urgently required and their discovery and development is the topic of intense investigation [3]. β-lactam/β-lactamase inhibitor combinations currently in clinical use include amoxicillin/clavulanic acid, ampicillin/sulbactam and piperacillin/tazobactam (TZP), with ceftolazone/tazobactam and ceftazidime/avibactam recently introduced into clinical use. TZP has broad spectrum antibacterial activity and is routinely used for intra-abdominal infections and febrile neutropenia [4, 5]. TZP usage has increased year on year in the UK, from just under 2.1% of all antibiotics prescribed in 2008-2009 to 3.6% in 2012-2013 [6]. During the 2-year period between April 2012 and March 2014 10.2% of bacteraemia causing *Escherichia coli* isolates in England tested for TZP susceptibility were resistant [7]. Resistance to TZP has been previously linked to AmpC hyperproduction, and the co-production of multiple β-lactamases, which also confer resistance to 3^rd^ generation cephalosporins [8]. Additionally, tazobactam is a poor inhibitor of metallo-β-lactamase enzymes, which cause resistance to TZP, alongside 3^rd^ generation cephalosporins and carbapenems [9–11].

A novel phenotype in *Klebsiella pneumoniae* and *E. coli* clinical isolates has emerged which has been classified as TZP-non-susceptible but susceptible to 3^rd^ generation cephalosporins and carbapenems [12–14], indicating a different resistance mechanism. While still relatively rare, one study in the United States found that the frequency of this phenotype was between 1.9% and 5.6% of *E. coli* and *K. pneumoniae* isolated from the bloodstream between 2011 and 2015, and specifically 4.1% of all *E. coli* over the study period [14]. The same study reported risk factors associated with the TZP-non-susceptible but 3^rd^ generation cephalosporin and carbapenem susceptible phenotype, including exposure to β-lactam/β-lactamase inhibitors and cephalosporins within the previous 30 days [14]. Resistance to TZP, but 3^rd^ generation cephalosporin and carbapenem susceptible, has been linked to hyperproduction of the β-lactamase *bla*_TEM_ which can hydrolyse piperacillin but not 3^rd^ generation cephalosporins. Hyperproduction is hypothesised to overcome the inhibitory activity of tazobactam, allowing the hydrolysis of piperacillin [15]. Mechanisms leading to hyperproduction include mutations in the promoter region of *bla*_TEM_, changing it from a weak promoter (*P3*) to a stronger promoter (*P4* or *P5*) [16] or a single point mutation further upstream resulting in the overlapping, stronger promoter *Pa/Pb* superseding the weaker P3 promoter [17], and increased production of *bla*_TEM_. Another such mechanism proposed to cause TZP-resistance but 3^rd^ generation cephalosporin and carbapenem susceptibility is increase in copy number of *bla*_TEM_ present in the chromosome [15, 18]. Gene amplification has been linked to the cause of temporary antibiotic resistance seen in a sub-population of bacteria and is known as heteroresistance. Heteroresistance is often lost after multiple generations in the absence of antibiotic selective pressure, due to the fitness cost imposed by the production of extra proteins as a result of amplification [19, 20]. While the mechanism of amplification of the *bla*_TEM_ is not well known, recent studies have found that the amplified *bla*_TEM_ has been co-located on a segment of DNA containing other antibiotic resistance genes, such as *aadA* and *sulI*, termed a genomic resistance module [15]. Amplification of *bla*_TEM_ leading to TZP resistance via β-lactamase hyperproduction has also been suggested to be mediated by the presence of the insertion sequence, IS*26* [18]. IS*26* is often linked with the movement of antibiotic resistance genes; for example a translocatable unit (TU) containing IS*26* has been shown to be able to excise from the transposon Tn*4352*B, which itself was located on a plasmid, between two IS*26*, leaving one in the plasmid [21, 22]. Following excision, the single IS*26* and antibiotic resistance gene(s) found between the two insertion sequences forms a circular TU, which then can insert into a plasmid via a conservative Tnp*26*-dependent but RecA-independent mechanism, Tnp26 replicative transposition or RecA-dependent homologous recombination, preferentially adjacent to another IS*26* insertion sequence [21–23].

Here, using a clonal pair of *E. coli* isolated from a single patient from two separate infection episodes which displayed within-patient evolution to TZP resistance, we sought to determine the mechanism of gene amplification resulting in hyperproduction of *bla*_TEM-1B_. We found that a TU was excised from a novel composite transposon flanked by IS*26* present in the chromosome, leading to gene amplification which did not carry any fitness cost. We were also able to replicate the evolutionary event leading to the excision in the TZP-susceptible isolate and captured the excised TU in a plasmid containing a copy of IS*26*.

## Materials and methods

### Bacterial isolates, media and antibiotics

Clinical isolates of *E. coli* isolated from blood cultures between 2010 and 2017 at the Royal Liverpool University Hospital (Liverpool, UK) which were found to be carbapenem and cephalosporin susceptible but TZP resistant using the disc diffusion method of antimicrobial susceptibility testing (AST) were initially identified from isolate records. Isolate records were then searched for a corresponding carbapenem, cephalosporin and TZP susceptible isolate, isolated in the same or a previous infection episode from the same patient. Using these criteria, we identified five paired clinical isolates of *E. coli*. All isolates had been stored at the time of blood culture isolation in glycerol broth at −80°C.

All isolates were grown on LB (Lennox) agar at 37°C for 18 hours followed by growth in LB (Lennox) broth, (LB, Sigma, UK), iso-sensitest (ISO, Oxoid, UK) or M9 (50% (v/v) M9 minimal salts (2x) (Gibco, ThermoFisher Scientific, USA), 0.4% D-glucose, 4mM magnesium sulphate (both Sigma, UK) and 0.05 mM calcium chloride (Millipore, USA)) at 37°C for 18 hours at 200 rpm.

Piperacillin, tazobactam (both Cayman Chemical, USA), gentamicin (GEN), and amoxicillin trihydrate:potassium clavulanate (4:1, AMC) were solubilised in molecular grade water (all Sigma, UK), while chloramphenicol (CHL) and tetracycline (TET) (both Sigma, UK) were solubilised in ethanol (VWR, USA) and ciprofloxacin (CIP) was solubilised in 0.1N hydrochloric acid solution (both Sigma, UK). All stock solutions of antibiotics were filter sterilised through a 0.22μm polyethersulfone filter unit (Millipore, USA). In all assays, unless stated, tazobactam was used at a consistent concentration of 4 μg/ml and the piperacillin concentration was altered.

### Restriction Enzyme Digestion

Restriction fragment length polymorphism (RFLP) analysis of 1 μg of 16S rRNA PCR amplicon from the 10 putative clonal isolates were digested with AlwNI, PpuHI and MslI (all New England Biolabs, USA) and 1 μg of long fragment genomic DNA extracts of 153964, 152025, 190693 and 169757 were digested with SpeI and MslI (both New England Biolabs, USA) for 1 hour at 37°C. Both RFLP digest reactions were incubated for 5 minutes at 80°C and immediately run on a 1% agarose gel. Enzyme digest of 500 ng of plasmid DNA extracted from 190693 and 169757 was performed with PpuMI and XhoI (both New England Biolabs, USA) and immediately run on a 1% agarose gel following incubation for 1 hour at 37°C.

### Antimicrobial susceptibility testing

Initial AST for cefpodoxime (CPD), cefoxitin (FOX), TZP, meropenem (MEM), CIP, cefotetan (CTT), amikacin (AMK), ertapenem (ETP), AMC, chloramphenicol (CHL) and ampicillin (AMP) was performed in clinic using the disk diffusion method, according to the CLSI guidelines for Antimicrobial Susceptibility Testing [24].

AST for TZP, GEN, CIP, CHL, AMC and TET were performed using the broth microdilution method for minimum inhibitory concentrations, performed in cation adjusted Mueller Hinton Broth (CA-MHB), following CLSI Guidelines. Efflux pump inhibition was performed using phenylalanine-arginine β-naphthylamide (PAβN) as a supplement in CA-MHB at a final concentration of 50μM.

### Nitrocefin assay

β-lactam hydrolysis was evaluated using a colorimetric nitrocefin assay. Cell lysates were obtained from triplicate cultures of 190693 and 169757 in LB, adjusted to an OD_600_ of 0.1. Cultures (10ml) were centrifuged at 14,000 g for 5 minutes, the supernatant discarded, and the pellet resuspended in 5ml phosphate buffered saline (PBS). The cultures were sonicated for three intervals of ten seconds, on ice, using a Soniprep 150 plus (MSE centrifuges, UK). The lysed cultures were centrifuged at 14,000 g for 5 minutes, and the supernatant taken as the culture lysate.

A total of 90μl of this lysate was then added to 10μl of 0.5mg/ml nitrocefin solution (Sigma, UK) in a 96 well microplate, in triplicate. The absorbance of the plate was read at 450nm every 20 seconds for 25 minutes, using a SPECTROstar OMEGA spectrophotometer (BMG lab systems).

### Whole genome sequencing and bioinformatics

Illumina MiSeq 2 × 250 bp short-read sequencing of long fragment DNA extractions from isolates 190693 and 169757, as well as adapter trimming of the sequencing reads, were provided by MicrobesNG (MicrobesNG, UK).

The same long fragment DNA extracts were processed using the SQK-LSK109 ligation and SQK-RBK103 barcoding kit and sequenced on an R9.4.1 flow cell with an Oxford Nanopore Technologies (ONT) MinION. Sequencing reads were basecalled during the sequencing run using MinKNOW, de-multiplexing and adapter trimming of the basecalled reads were performed using Porechop (v0.2.4) and finally sequencing reads were filtered for a quality score of 10 via Filtlong (v0.2.0).

Both Illumina short-read and MinION long-read sequences were assembled using Unicycler (v0.4.7 [25]), with the quality of the assembly assessed using QUAST (v5.0.2 [26]), annotated using Prokka (v1.14.0 [27]) and visualised using Bandage (v0.8.1 [28]).

Sequence type and serotype of both 190693 and 169757 were determined using Multi-Locus Sequence Typing (MLST, v2.0.4 [29]) and SerotypeFinder (v2.0.1 [30]), respectively. The relatedness of the two genomes were compared using MUMmer (v3.23 [31]) and the average nucleotide identity (ANI) was calculated using OrthoANI (v0.93.1 [32]). Presence of acquired antimicrobial resistance genes within the two genomes were assessed using ResFinder with minimum threshold of 90% and a minimum length of 60% (v3.2 [33]) and segments of the two genomes were characterised using SnapGene^®^ software (from GSL Biotech; available at snapgene.com). Finally, plasmid replicons were identified using PlasmidFinder (v2.0.1) [34].

### Competent cell preparation

The TZP-susceptible isolate was made competent according to Chung *et al.* [35].

### Quantitative PCR

Changes in gene copy number of *bla*_TEM-1B_, *bla*_OXA-1_, *aac*(3)-*lla*, *aac*(6’)-*lb-cr*, *tet*(D) were calculated via qPCR, using the ΔΔCT method for relative quantitation of these genes against the single copy *uidA* housekeeping gene.

Each qPCR reaction contained 6.25μl QuantiTect^®^ SYBR Green PCR buffer (Qiagen, UK), 0.4 μM forward and reverse primers (Table S1), 1ng of extracted DNA, and molecular grade water to a final volume of 12.5μl. Reactions were processed using a Rotor-Gene Q (Qiagen, Germany), using the following protocol; an initial denaturation step of 95°C for 5 minutes, followed by 40 cycles of; DNA denaturation at 95°C for 10 seconds, primer annealing at 58°C for 30 seconds, and primer extension at 72°C for 10 seconds with fluorescence monitored in the FAM channel. HRM analysis was carried out over a temperature range of 75°C to 90°C, increased in 0.1°C increments, in order to confirm specific amplification. Fluorescence thresholds were set manually for calling Ct values, at 5% of the difference between baseline and maximum fluorescence.

The mean qPCR Ct value for the *uidA* gene from each strain was taken using four replicate qPCR reactions, and the ΔΔCT method was utilised to determine fold change using quadruplicate qPCR reactions for each AMR gene.

### In vitro evolution of susceptible isolate

The clinical isolate 190693 (TZP-susceptible isolate) and 190693 transformed with pHSG396:IS*26* were subcultured into 10 ml LB and 10 ml LB plus 35 μg/ml chloramphenicol, respectively, and incubated at 37°C for 18 hours at 200 rpm. Following incubation 10 μl of 190693 was subcultured into 10 ml LB and 10 ml LB plus 8/4 μg/ml TZP and 10 μl of 190693 with pHSG396:IS*26* was subcultured into 10 ml LB plus 35 μg/ml chloramphenicol and 10 ml LB plus 35 μg/ml chloramphenicol and 8/4 μg/ml TZP and incubated at 37°C for 24 hours at 200 rpm. Genomic DNA from each of the four cultures were extracted for qPCR following the protocol described above, however triplicate biological replicates were used instead of quadruplicate qPCR reactions

### Translocatable unit capture

The TZP-susceptible isolate 190693 transformed with pHSG396:IS*26* was grown in the presence of TZP and chloramphenicol as previously stated. Following selection, the culture was serially diluted 1/10 in PBS down to 10^−7^ dilution and 50 μl of each dilution was plated out on to LB agar supplemented with 35 μg/ml chloramphenicol and 16/4 μg/ml TZP. Five single colonies were selected and subcultured into 10 ml LB plus 35 μg/ml chloramphenicol and 16/4 μg/ml TZP for 18 hours at 37°C and 200 rpm and the plasmid extracted following the protocol above. The purified plasmids were transformed into NEB^®^ 5-alpha competent *E. coli* following the protocol in the supplementary material and plated out on to LB agar supplemented with 35 μg/ml chloramphenicol and 16/4 μg/ml TZP and incubated at 37°C for 18 hours. A single colony from each transformation was subcultured into 10 ml LB supplemented with 35 μg/ml chloramphenicol and 16/4 μg/ml TZP and incubated at 37°C, 200 rpm for 18 hours and the plasmid extracted following the protocol in the supplementary material. The initial pHSG396:IS*26* plasmid extract and pHSG396:IS*26* plasmid selected in TZP and extracted from NEB^®^ 5-alpha *E. coli* were digested with XhoI (New England Biolabs, US) and EcoRI for 1 hour at 37°C, followed by a 20 minute incubation at 65°C and run on a 1% agarose gel.

### Competitive fitness

The relative fitness of 169757 (TZP-resistant) and 190693 (TZP-susceptible) grown in the presence of 8/4 μg/ml TZP, compared to 190693 and 190693 grown in the absence of TZP, were assessed comparatively in LB, ISO and M9. Each culture was diluted to an OD_600_ of 0.1 in the respective media, then further diluted 1/1000 in the same media and 150 μl of each diluted culture added to a flat bottom, 96 well microtitre plate in duplicate as well as 150 μl of the media as a negative control. The 96 well plate was incubated at 37°C and the OD_600_ of each well was measured with 100 flashes every 10 minutes over 24 hours, with orbital shaking at 200 rpm between readings, using a Clariostar Plus microplate reader (BMG Labtech, Germany). The relative fitness compared to either 190693 or 190693 grown in the absence of TZP between absorbance values 0.2 and 0.08 and a minimum R value of 0.9799 was estimated using BAT version 2.1 [36].

### Statistical analysis

Statistical analysis of comparison for qPCR of the antibiotic resistance genes on the RM of 190693 and 169757 was performed using the 2way ANOVA test. Statistical analysis of comparison for qPCR of the antibiotic resistance genes on the RM of 190693 grown in the presence or absence of TZP, with and without pHSG396:IS*26* was performed using the Ordinary One-Way ANOVA with Uncorrected Fisher LSD test. Statistical analysis of relative fitness of 169757 and 190693 grown in the presence of TZP was performed using Ordinary One-Way ANOVA with Uncorrected Fisher LSD test. All statistical tests were performed using GraphPad Prism version 8.2.1.

## Results

### Identification of clonal isolates

Initially, we identified five isolates in the collection of TZP-resistant, 3^rd^ generation cephalosporin and carbapenem susceptible *E. coli* from blood cultures at the Royal Liverpool University Hospital which had a corresponding TZP-susceptible isolate from the same or previous infection episode, and therefore could have evolved to become TZP-resistant within a patient. RFLPs of the 16S rRNA amplicons from the five pairs of isolates indicated that three pairs of TZP-susceptible/TZP-resistant clinical isolates had identical digestion patterns (Fig. S1A). Two of these three pairs of isolates had an identical resistance profile generated during routine disk-based susceptibility testing, aside from TZP (Table S2). RFLPs of genomic DNA identified one pair of isolates with identical banding patterns indicating clonality; 190693 (TZP-susceptible) and 169757 (TZP-resistant) which were isolated from different infection episodes from the same patient approximately 3 months apart (Fig. S1B). During the first infection episode, the TZP-susceptible *E. coli* was isolated and the patient was initially treated with a five day course of TZP, followed by a seven day course of TZP with teicoplanin and then a third seven day course of TZP although a second blood culture was found to be negative. A second infection episode occurred approximately 6-7 weeks after the final course of TZP was completed, and again the patient was treated initially with TZP until the TZP-resistant *E. coli* was isolated, when the treatment was changed to meropenem. Putative clonality of these two isolates was confirmed with whole genome sequencing; both isolates were identified as serotype H30 O86, sequence type 8 and had an ANI of 100%, with 36 single nucleotide polymorphisms difference between the two isolates.

### Confirmation of TZP susceptibility/resistance and mechanism of resistance

We determined the MIC of the pair of isolates and verified that TZP-susceptible isolate was susceptible to TZP (2-4/4 μg/ml) and TZP-resistant isolate was resistant to TZP (64/4 μg/ml) according to EUCAST clinical breakpoints [37] (Table 1). Using the efflux pump inhibitor PAβN as a supplement in the MIC assay, we were able to rule out overexpression of efflux pumps as a possible mechanism of resistance as there was less than a 4-fold reduction [15] in MIC of both the TZP-susceptible (2/4 μg/ml) and TZP-resistant isolates (32/4 μg/ml, Table 1). Whole genome sequencing revealed no differences in the predicted resistance genes present in the genome between the TZP-susceptible and TZP-resistant isolate (Table 2) and no mutations in the promoter region of any of the BLs present within the genome (*bla*_TEM-1B_, *bla*_OXA-1_ or *ampC*) of the TZP-resistant isolate. We confirmed that the TZP-resistant isolate hyperproduced a BL due to the significant increase in nitrocefin hydrolysis compared to the TZP-sensitive isolate (P value = <0.0001, Fig. 1A).

**Table 1:**
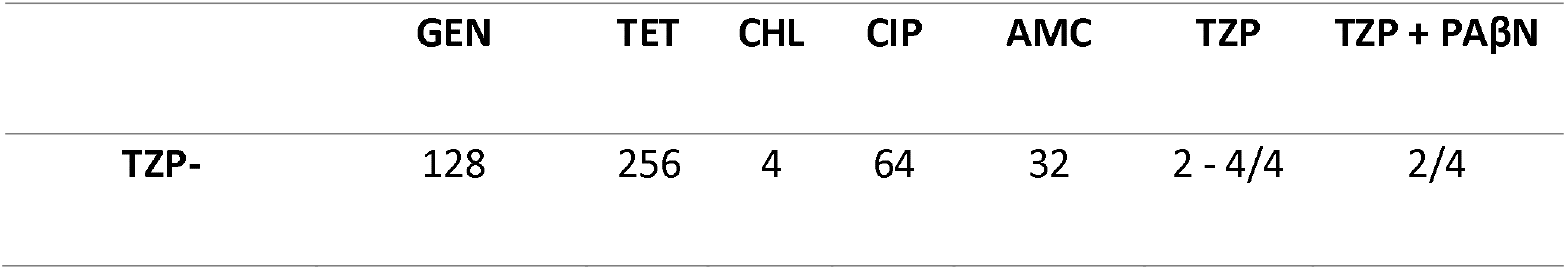
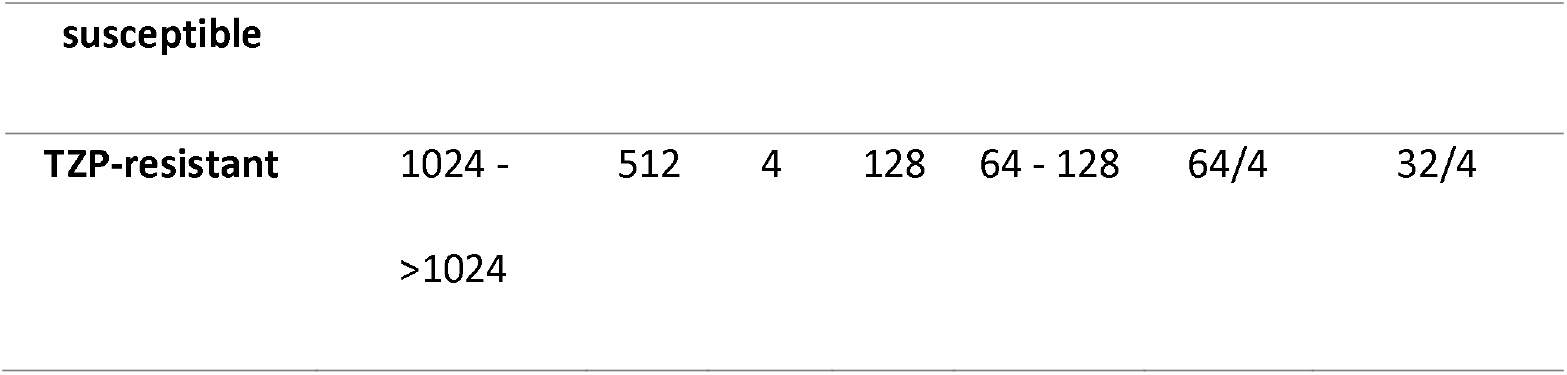
Minimum inhibitory concentrations of gentamicin (GEN), tetracycline (TET), chloramphenicol (CHL), ciprofloxacin (CIP), amoxicillin/clavulanic acid (AMC) and piperacillin/tazobactam (TZP) (with and without PAβN) towards the TZP-susceptible and TZP-resistant isolates

**Table 2:** Predicted antimicrobial resistance genes by ResFinder found on the genome of the TZP-susceptible and TZP-resistant isolates and their position in genome. *The resistance gene catB3 was predicted by ResFinder to be present with 69.8% length, but both isolates were phenotypically chloramphenicol susceptible

| Antimicrobial Resistance Gene | TZP-susceptible isolate | TZP-resistant isolate |
| --- | --- | --- |
| *bla<sub>OXA-1</sub>* | Chromosome | Translocatable unit |
| *bla<sub>TEM-1B</sub>* | Chromosome | Translocatable unit |
| *aac(3)-IIa* | Chromosome | Translocatable unit |
| *aac(6')-Ib-cr* | Chromosome | Translocatable unit |
| *aadA1* | Chromosome | Chromosome |
| *aph(3'')-Ib* | Chromosome | Chromosome |
| *aph(6)-Id* | Chromosome | Chromosome |
| *tet(D)* | Chromosome | Translocatable unit |
| *dfrA1* | Chromosome | Chromosome |
| *sul2* | Chromosome | Chromosome |
| *mdf(A)* | Chromosome | Chromosome |

| *catB3* * | Chromosome | Translocatable unit |
| --- | --- | --- |

**Figure 1:**
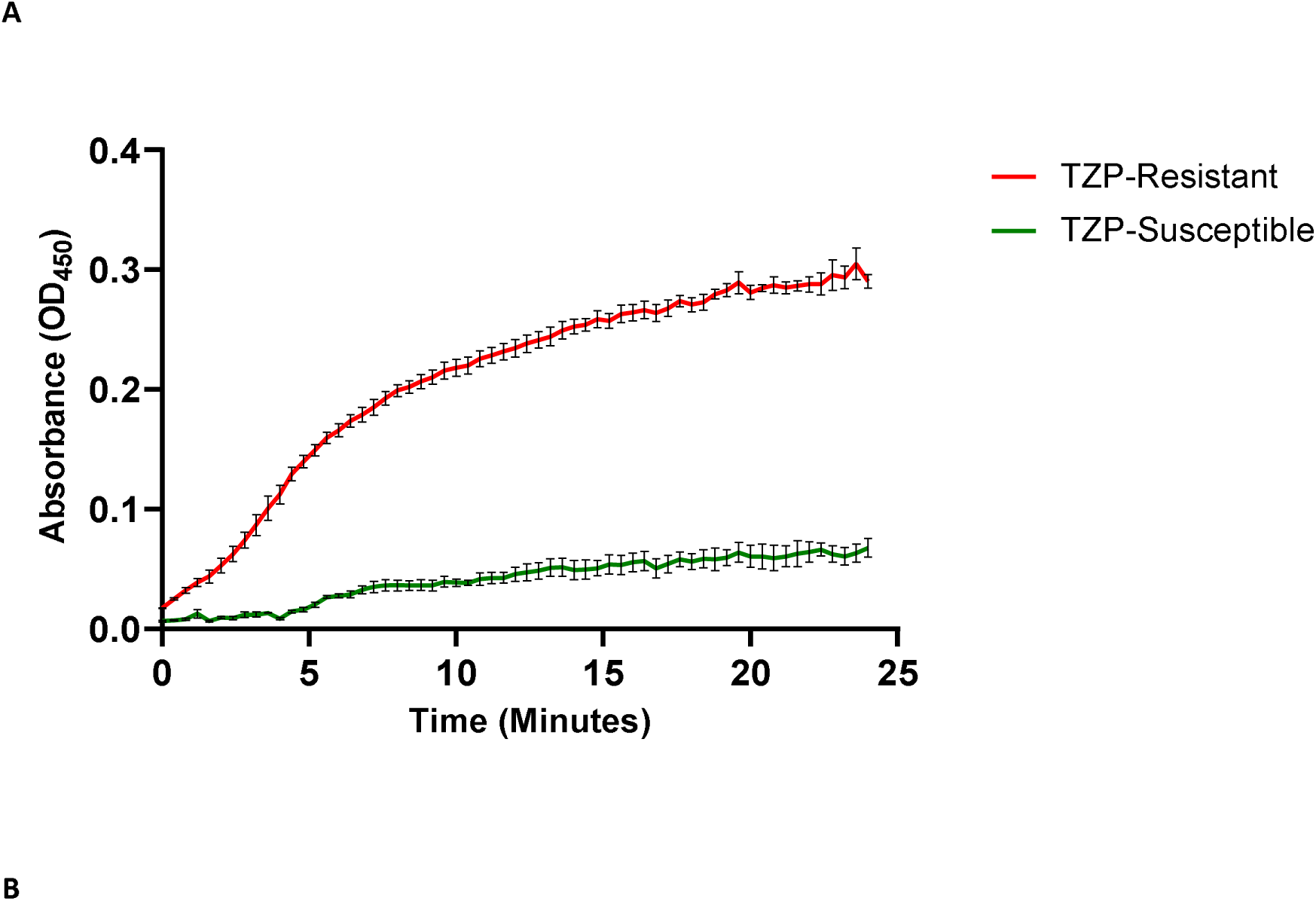
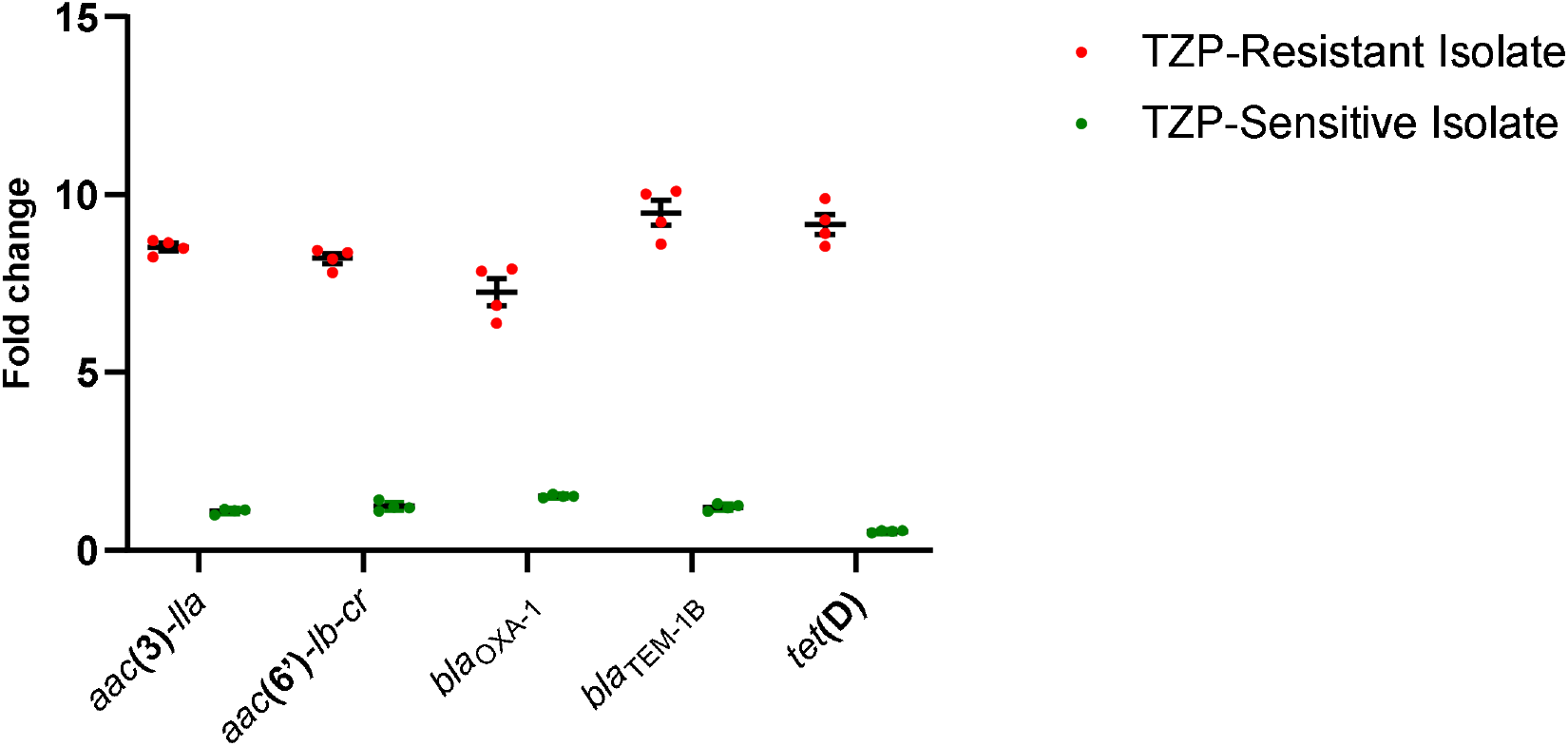
**A**) Increase in nitrocefin hydrolysis by the TZP-resistant isolate compared to the TZP-susceptible, isolate as measured at OD_450_, due to hyperproduction of a BLS in the TZP-resistant isolate. **B**) Comparison of the fold change in copy number of the antimicrobial resistance genes present on the composite transposon of the TZP-susceptible isolate/translocatable unit of the TZP-susceptible isolate as assessed by qPCR compared to the housekeeping gene *uidA*

### Hybrid assembly and comparison of the genomes of the TZP-susceptible and TZP-resistant isolates

Using a hybrid assembly of ONT long and Illumina short sequencing reads, we were able to complete the genome of the TZP-susceptible isolate, which was found to be 5151952 bp in length, with a GC content of 50.64% and did not contain any plasmids (Fig. S4A). In contrast, we unable to complete the genome of the TZP-resistant isolate as a 530bp segment remained unresolved, and a complete, low copy number (2.59x) 106637 bp plasmid containing an IncFII replicon (Fig. S4B) was detected. A complete, smaller (10899 bp) circular DNA molecule was also found to be present in the TZP-resistant isolate, at a copy number of 8.51x, however this small circular DNA molecule did not contain a plasmid replicon (Fig. S4B). The large plasmid did not to contain any predicted antimicrobial or metal resistance genes, but did contain three bacteriocins, both colicin B and M (with cognate immunity proteins) and linocin. Comparison of the predicted resistance genes present on the chromosome of the TZP-susceptible and TZP-resistant isolates highlighted that *bla*_TEM-1B_, *bla*_OXA-1_, *aac*(3)-*lla*, *aac*(6’)-*lb-cr*, *tet*(D) and *cat*B3 were missing from the assembled chromosome of the TZP-resistant isolate (Table 2). Characterisation of the small circular DNA molecule found that it contained these missing resistance genes, as well as several putative transposable elements including three copies of IS*26* (Table 2), and aligned exactly to the chromosome of the TZP-susceptible isolate and was no longer present in the chromosome of the TZP-resistant isolate. The predicted *cat*B3 resistant gene was truncated to 69.8% length and therefore unlikely to be functional, which was confirmed as both the TZP-susceptible and TZP-resistant isolates were sensitive to chloramphenicol according to EUCAST guidelines (Table 1). Further analysis of the TZP-susceptible genome uncovered that the circular DNA molecule from the TZP-resistant isolate aligned with 100% identity to a novel integrated composite transposon flanked by two copies of IS*26* in the same orientation, which we subsequently registered as Tn*6762* via the transposon registry [38], however the circular DNA molecule only contained one IS*26*. The antibiotic resistance gene *bla*_TEM-1B_, *bla*_OXA-1_, *aac*(3)-*lla*, *aac*(6’)-*lb-cr* and *tet*(D) were identified to be present on Tn*6762* (Fig. 2A). This suggested that a translocatable unit (TU) [21–23], containing one IS*26* and the antibiotic resistance genes, was excised from the chromosomally located Tn*6762* while the other IS*26* stayed in the chromosome (Fig. 2A). Interestingly, we also found that the tetracycline resistance regulator, *tetR*, on the TU had been disrupted 52 bp from the end of the tetR due to the excision which overlaps the start of the copy of IS*26* remaining in the chromosome. Following excision, a copy of IS*26* is present at the start of the TU which then connects to the end of the TU, containing the disrupted *tetR*, forming a circular DNA molecule. As the two copies of IS*26* are identical and in the same orientation, and therefore containing the same 52 bp bases missing from *tetR*, tetR was reformed when the TU circularised completing the 657 bp gene.

**Figure 2:**
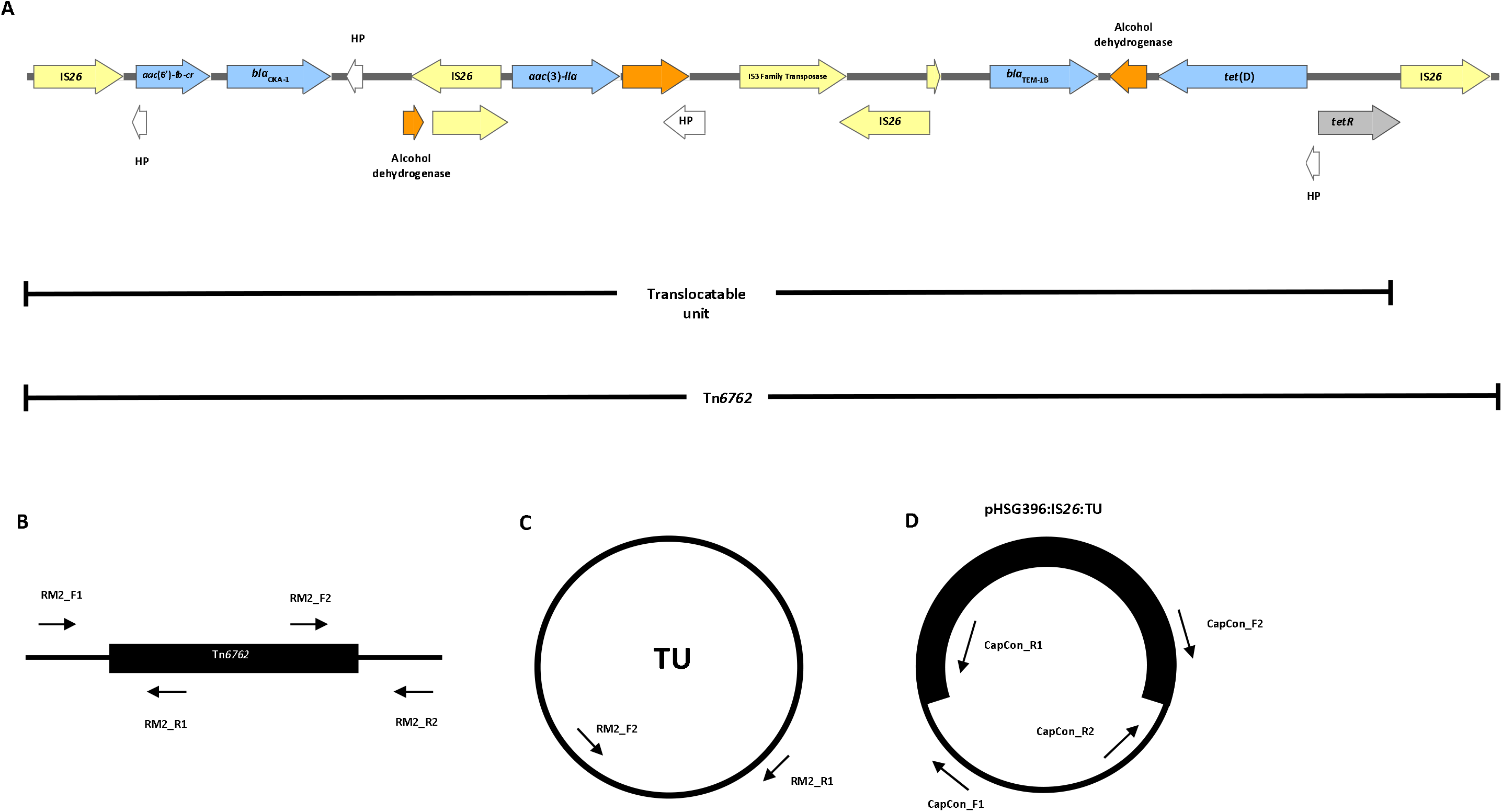
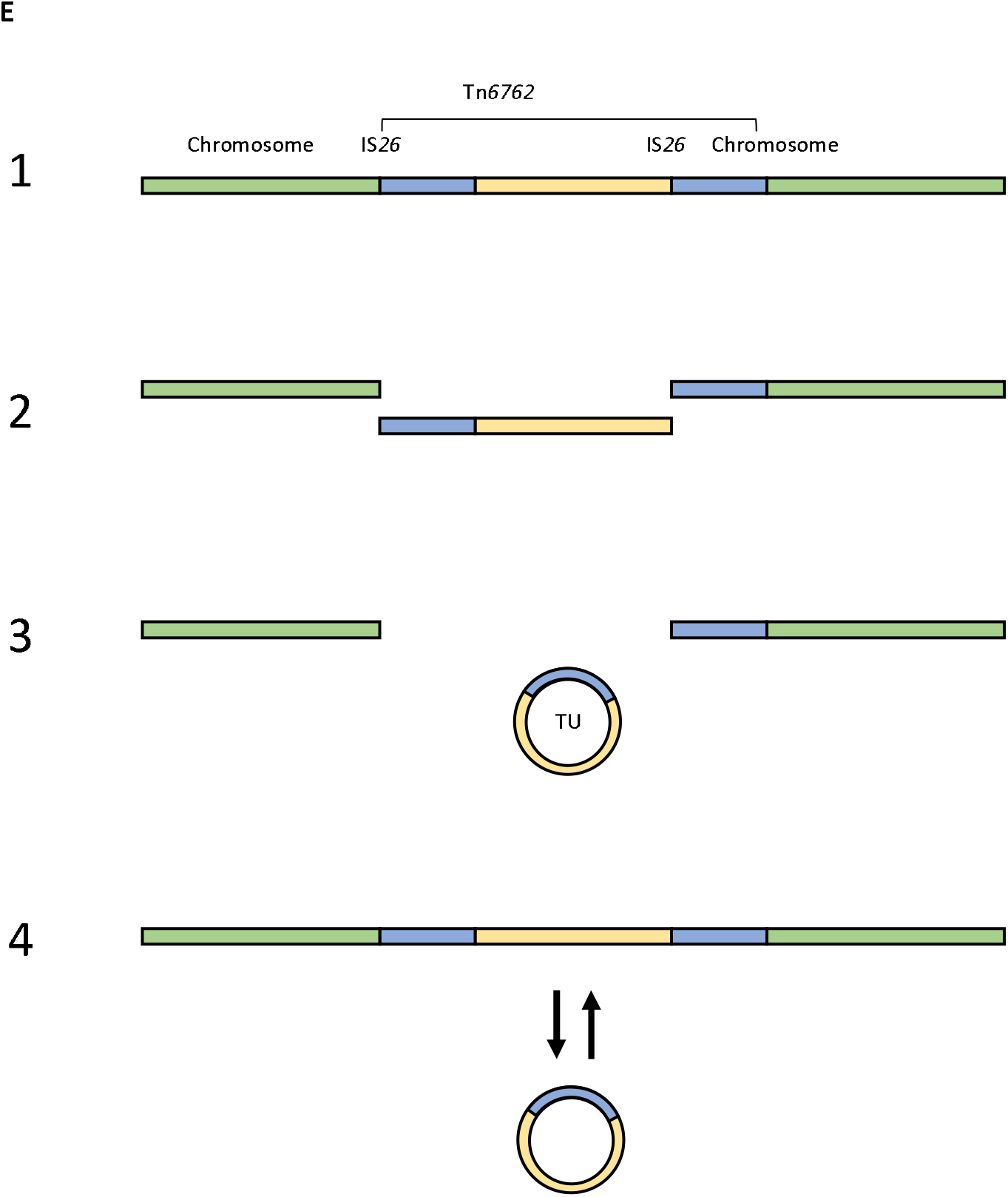
Schematic showing **A**) the characterisation of Tn*6762* (HP = hypothetical protein) and the position of the primer pairs to detect **B**) the junctions of the Tn*6762* in the chromosome, **C**) the presence of the TU and **D**) the junctions of the insertion of the TU into pHSG396:IS*26*. **E**) Schematic of the proposed mechanism of leading to hyperproduction of *bla*_TEM-1B_; **1**) a composite transposon, Tn*6762*, is present on the chromosome flanked by two copies of IS*26*. Due to selective pressure from TZP, **2**) Tn*6762* is excised from the chromosome, **3**) which then forms a translocatable unit (TU) which **4**) re-inserts and excises from the chromosome adjacent to the chromosomally located IS*26*, increasing the copy number of *bla*_TEM-1B_

### Confirmation of the presence of Tn6762 and the translocatable unit

Firstly, we confirmed the amplification of the resistance genes found on the TU in the TZP-resistant isolate and Tn*6762* in the chromosome of the TZP-susceptible isolate. Comparing the fold change in copy number of each resistant gene to the housekeeping gene *uidA*, we found that each resistance gene on the TU increased in copy number in the TZP-resistant isolate compared to the TZP-susceptible isolate (Fig. 1B, P value = <0.0001). The increase in copy number of the resistance genes found on the TU in the TZP-resistant isolate also corresponded to an increase in MIC of all antimicrobials that the genes confer resistance to (Table 1), further confirming the amplification of the entire TU. By PCR of the left and right junctions of the chromosomally located Tn*6762*, with one primer specific for Tn*6762* and one for chromosome either before or after IS*26* (Fig. 2B), we were able to confirm that Tn*6762* was present in the chromosome of both the TZP-susceptible isolate and TZP-resistant isolate by yielding the expected 1640 bp and 2402 bp products, respectively (Fig. S2). Using primers that would only yield a 1942 bp product if the circular TU was present (Fig. 2C), we were able to detect the presence of the TU in the TZP-resistant isolate and absence in the TZP-susceptible isolate (Fig. S2). This suggests that Tn*6762* has been excised and existed as a circular TU, as well as in the chromosome of the TZP-resistant isolate.

### Replication of excision of the translocatable unit

Completed assembly of the large plasmid present in the TZP-resistant isolate showed that the plasmid contained a single copy of IS*26*. We sought to determine whether the presence of another mobile IS*26* and/or the use of TZP induced the excision of the TU by replicating the evolutionary event that led to the TZP-sensitive isolate becoming resistant to TZP *in vitro*. We found that there was a significant increase in copy number of all the resistance genes present on Tn*6762*, relative to the housekeeping gene *uidA*, following exposure to 2 × MIC of TZP (8/4 μg/ml) while there was no evidence of amplification in the same isolate grown in the absence of TZP (P value = 0.0012). There was also an increase in copy number when the isolate containing the pHSG396:IS*26* plasmid was exposed to TZP but, again, no increase in copy number when TZP was absent (P value = <0.0001), therefore TZP can either select for the maintenance of the excised TU or induce the excision event, leading to an increase in copy number of the genes present on Tn*6762*. In contrast, there was no significant difference in gene copy number between the TZP-susceptible with and without the pHSG396:IS*26* plasmid grown in the absence of TZP (P value = 0.5064), underlining that the presence of an extra chromosomal IS*26* does not induce excision of the TU from Tn*6762*.

### Capture of the translocatable unit

We sought to capture the excised TU, and therefore observe the excision and insertion events, using a pHSG396 plasmid containing IS*26* transformed into the TZP-sensitive isolate in the same orientation relative to the origin of replication as found on Tn*6762* and selected for TZP-resistant derivatives by growing the isolate in the presence of TZP. We detected the insertion of a >10kb fragment into the pHSG396:IS*26* after TZP selection following digestion with XhoI and EcoRI (Fig. S3A). Insertion of the TU from the TZP-susceptible chromosome was confirmed through PCR amplification across the two newly formed junctions on the pHSG396:IS*26*, with one primer specific for pHSG396 and one for either *aac*(6’)-*lb-cr* (left) or *tet*(D) (right) on the TU for each junction yielding the expected 1458 bp (left) and 1385 bp (right) products (Fig. 2D), consistent with the insertion of the TU adjacent to IS*26* in the pHSG396 and reforming Tn*6762* in pHSG396 (Fig. S3B).

### Fitness effect of extensive amplification

Hyperproduction of a protein could result in a fitness cost to the cell due to the increased metabolic activity. Yet, we found that the amplification of the TU and carriage of the large plasmid in the TZP-resistant isolate did not result in a significant change in fitness compared to the TZP-susceptible isolate in LB (P value = 0.9968), ISO (P value = 0.2836) and M9 (P value = 0.2204, Fig. 4). We also assessed the relative fitness of the TZP-susceptible isolate following exposure to TZP (resulting in the increase in copy number of the resistance genes present on the RM (Fig. 3)) compared to the TZP-susceptible grown in the absence of TZP (which did not result in an increase in copy number (Fig. 3)). Again, we found no significant change in fitness in LB (P value = 0.8047), ISO (P value = 0.1242) and M9 (P value = 0.2803, Fig. 4).

**Figure 3:**
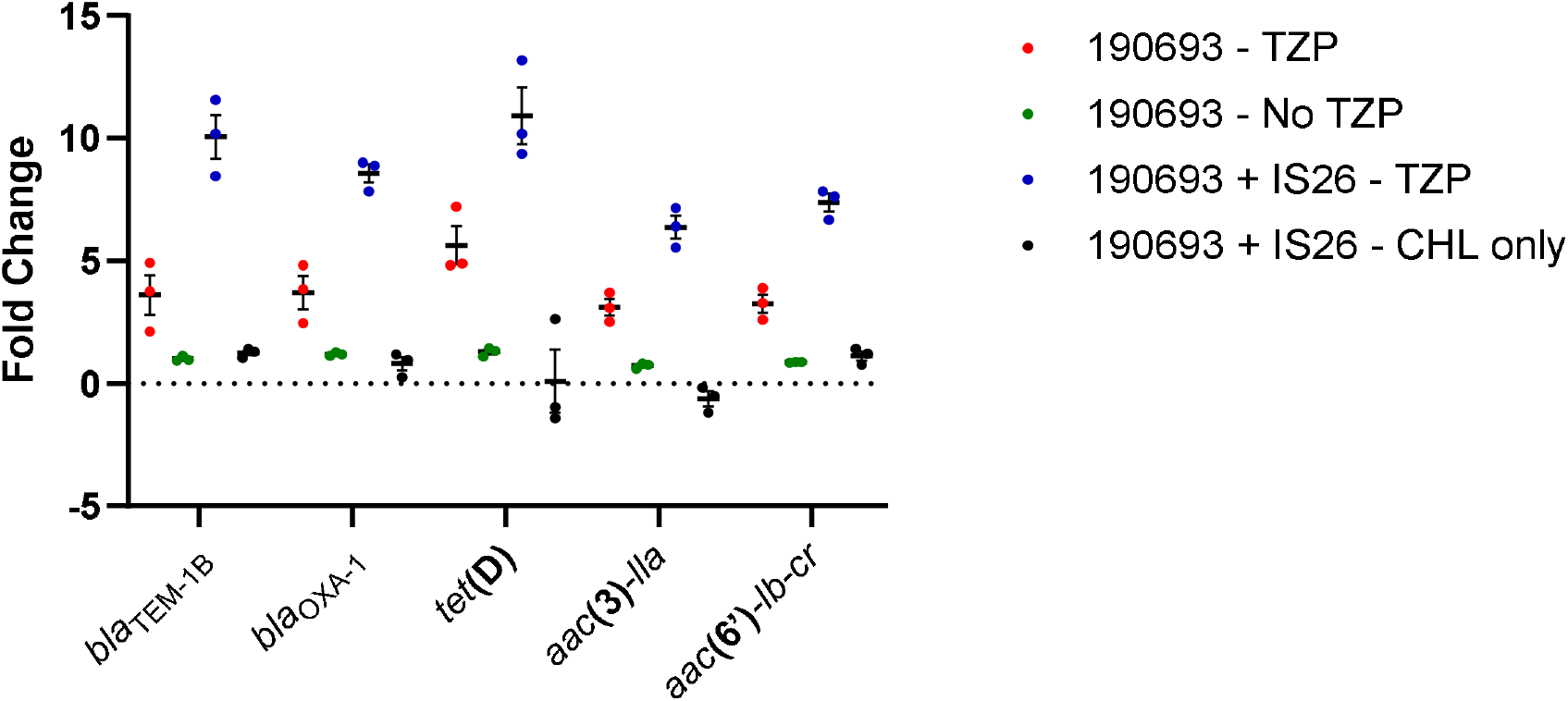
Fold change in copy number of all the antimicrobial resistance genes found on the composite transposon compared to the housekeeping gene *uidA* following growth of the TZP-susceptible isolate in the absence of antibiotics, TZP-susceptible isolate in the presence of 8/4 μg/ml TZP, TZP-susceptible isolate transformed with pHSG396 plasmid containing IS*26* in the presence of 8/4 μg/ml TZP and 35 μg/ml chloramphenicol and TZP-susceptible isolate transformed with pHSG396 plasmid containing IS*26* in the presence of 35 μg/ml chloramphenicol only

**Figure 4:**
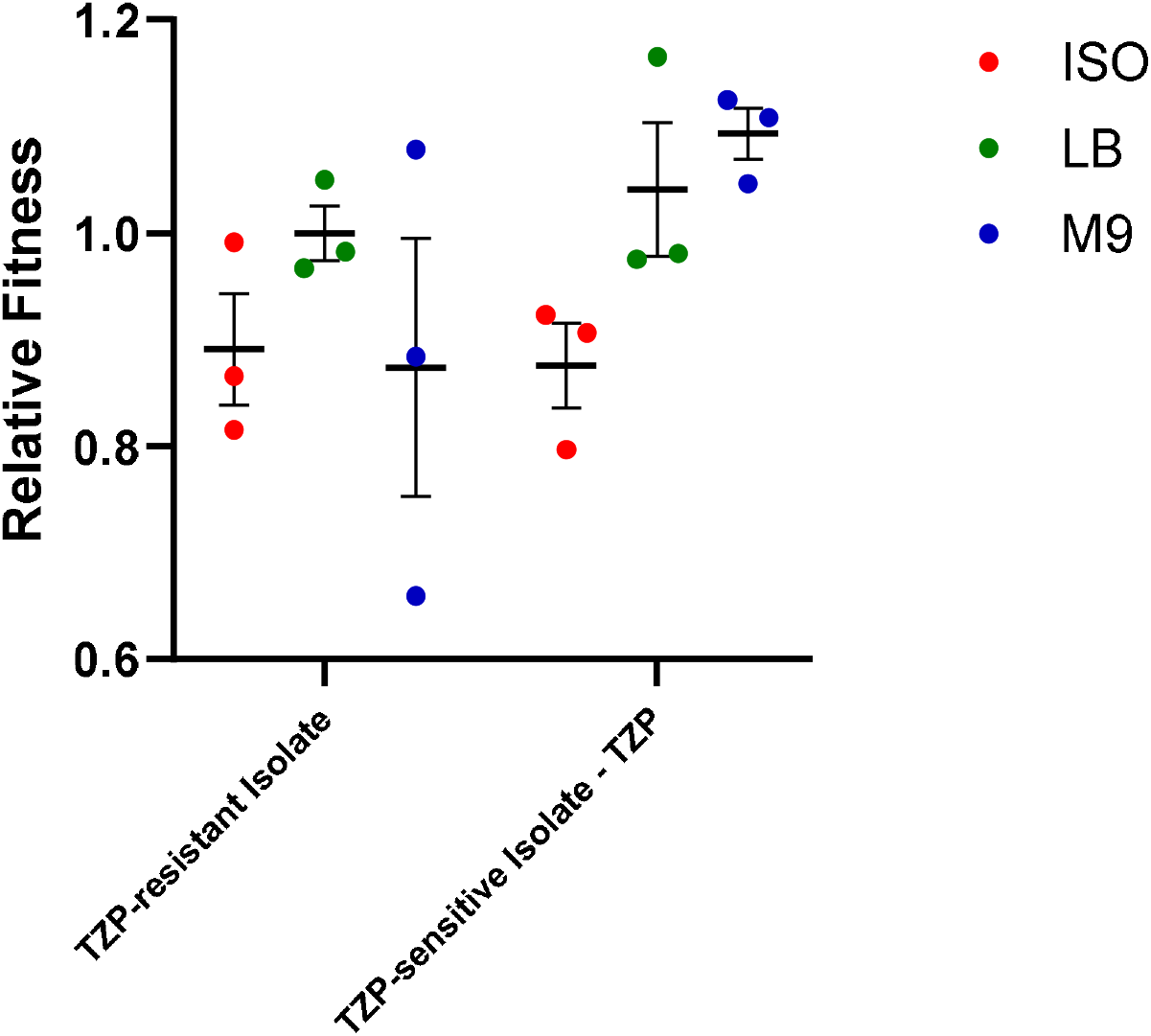
Relative fitness of the TZP-resistant isolate compared to the TZP-susceptible isolate and the TZP-susceptible isolate grown in the presence of TZP compared to the TZP-susceptible isolate grown in the absence of TZP, assessed comparatively in LB, ISO and M9

## Discussion

Tazobactam is able to inhibit the activity of class A β-lactamases [39], and therefore the presence of *bla*_TEM-1_ within the genome of an *E. coli* isolate should not result in resistance to TZP. However, two studies have linked amplification of *bla*_TEM_, and therefore hyperproduction of the BL, with this phenotype [15, 18], with one linking amplification and the presence of IS*26* [18]. However, the exact mechanism of amplification/hyperproduction has remained elusive. Due to the emergence of a novel TZP-resistant but 3^rd^ generation cephalosporins and carbapenems susceptible *E. coli* and *K. pneumoniae* phenotype [12–14], as well as the increasing reliance on TZP as an empirical treatment [6] and the recent interest in the use of TZP as a carbapenem sparing treatment for ESBL infections [40, 41], it is of growing importance to understand the mechanism of resistance. In this study we had the unique opportunity to compare a pair of clonal isolates which have evolved within a patient to become TZP-resistant but remain cephalosporin/carbapenem sensitive, allowing us to build on recent studies and identify the mechanism of IS*26*-mediated amplification of *bla*_TEM-1B_ which leads to TZP-resistance.

We found multiple antibiotic resistance genes, including *bla*_TEM-1B_, co-located on a novel IS*26* composite transposon Tn*6762* on both the TZP-resistant and TZP-susceptible isolate. This is the first time that the resistance genes *tet*D and *bla*_TEM-1B_ have found to be present on the same transposable element and a result of insertion of several transposons into the same location, demonstrated by the presence of several insertion sequences and transposable elements on Tn*6762*. Amplification of *bla*_TEM-1B_ is achieved when Tn*6762* is excised from the chromosome forming a TU, evidenced by the hybrid assembly of the TZP-resistant isolate, increase in copy number of the antibiotic resistance genes present on Tn*6762* and the capture of the TU in pHSG396:IS*26*. Precise excision and formation of a TU containing the antibiotic resistance gene aphA1a from a Tn*4352*B transposon present in a plasmid has been demonstrated before as a mechanism of movement of antibiotic resistance gene [21, 22, 42], this is the first time it has been shown to directly result in gene amplification resulting in resistance. We found no evidence from the whole genome sequencing of the TZP-resistant isolate of insertion of the TU anywhere else in the chromosome, except for a gap in sequencing where Tn*6762* was originally situated in the TZP-susceptible isolate adjacent to an IS*26*. Through PCR of the left and right junctions of the gap in sequencing, we confirmed that Tn*6762* was still present at this location in the chromosome of the TZP-resistant isolate and therefore existed both as Tn*6762* and TU, which the hybrid assembly was unable to resolve. We also found no evidence of a tandem repeat of the TU in the chromosome of the TZP-resistant isolate and only evidence of the TU through hybrid assembly of the genome and confirmation by PCR. Using the chloramphenicol-resistant pUC vector, pHSG396, containing a copy of IS*26* from the TZP-susceptible chromosome, we were able to replicate the evolutionary event and capture a single copy of the excised TU adjacent to the IS*26* copy in the plasmid, providing evidence of the insertion event, but again there was no evidence of tandem repeats of the TU in the pHSG396:IS*26* plasmid. Therefore, the TU preferentially re-inserts into the chromosome adjacent to the chromosomally located IS*26* via a conservative Tnp26-dependent but RecA-independent mechanism, Tnp26 replicative transposition or RecA-dependent homologous recombination [22]. While this mechanism has previously been demonstrated in terms of movement of antibiotic resistance genes [21, 22], this is the first time this mechanism has been shown to cause gene amplification resulting in antibiotic resistance in a clinical isolate, be able to replicate the evolutionary event resulting in amplification *in vitro* and show that use of TZP either selects for or induces this phenotype. This mechanism is of concern as IS*26* has been associated with the transfer of *bla*_NDM-1_ in a recent nosocomial outbreak in Germany [42], and other carbapenemases [43] and therefore represents a risk of clinical resistance to any carbapenemase inhibitor currently in development [44] which needs to be investigated further. The method of capture of the TU used in this study can be used to investigate whether the same mechanism will result in amplification to other β-lactam/β-lactamase inhibitors, as well as to further confirm the role of TZP in the induction of excision of the TU, and therefore gene amplification.

The absence of insertion of the TU into the plasmid in the TZP-resistant isolate was notable. After determining the direction of IS*26* in relation to the origin of replication in both the chromosome of the TZP-susceptible isolate and the plasmid, IS*26* was in the reverse direction in the plasmid but in the forward direction on both the TU and the chromosome. The orientation of IS*26* may have bearing on whether insertion of the TU can occur, as previous studies on the insertion of a TU via a conservative mechanism found that both IS*26* were in direct orientation [21–23], however this needs further investigation to confirm whether the direction of the IS*26* is critical for insertion of the TU.

Hansen *et al.* 2019 associated amplification of *bla*_TEM-1_ in *E. coli* clinical isolate with a significant fitness cost [18];Fitness of this isolate was compared to other unrelated clinical isolates of *E. coli* which hyperproduced *Bla*_TEM-1_ due to promoter mutations, rather than the same isolate with and without amplification. This approach can lead to over- or under-estimation of fitness cost, as genetic background of the isolate can have an impact on the overall fitness affect, as can the environment fitness is assessed in [45, 46]. Adler *et al.* 2014, however, identified a fitness cost associated with IS*26*-mediated amplification of an antibiotic resistance cassette from a plasmid in all lineages tested [20]. In this study, we were able to compare the fitness of the paired clinical isolate and *in vitro* evolved isolates with and without amplification of the chromosomally located composite transposon and found that there was no significant difference in fitness cost, despite amplification of a >10kb region with multiple functionally transcribed genes and, in terms of the TZP-resistant clinical isolate, has acquired a large plasmid. If this lack of fitness cost is translated into a physiological environment, it may result in the TZP-resistant phenotype persisting. While there was no observed fitness effect of amplification on this single isolate, the effect of amplification on bacterial fitness needs to be extensively investigated as it may not be a global phenomenon.

## Conclusions

Resistance to the β-lactam/β-lactamase inhibitor TZP can be the result of hyperproduction of the β-lactamase *bla*_TEM-1B_ due to the IS*26*-associated excision and circularisation of a translocatable unit containing *bla*_TEM-1B_ from the chromosome either selected by or in response to exposure to TZP. In this clinical isolate and an *in vitro* evolved isolate, we found that there was no effect on fitness due to the amplification and subsequent carriage of high numbers of the TU. This mechanism of amplification, and the subsequent hyperproduction, of *bla*_TEM-1B_ is an important consideration if treatment failure involving TZP occurs, as well as other β-lactam/β-lactamase inhibitor combinations, and when using genomic data to predict resistance/susceptibility to β-lactam/β-lactamase inhibitor combinations.

## Author contributions

ATMH and TE conceptualised the study. JM, PR, CMP, CC, JvA, and AH collated isolate metadata, clinical antimicrobial susceptibility data and patient treatment data. ATMH, AJF, ERA, APR and TE contributed to the experimental design and data analysis. ATMH, AJF and TE contributed to carrying out the experiments. ATMH and TE wrote the first draft of the manuscript, which was then edited and approved all authors by all authors.

## Data availability

Unicycler hybrid assemblies of the two clonal isolates of *E. coli* were submitted to GenBank under the BioProject PRJNA607545. Accession number CP048934 corresponds to TZP-susceptible isolate (190693) and accession number JAAKGF000000000 corresponds to TZP-resistant isolate (169757).

## Disclaimer

This report is independent research funded by the Department of Health and Social Care. The views expressed in this publication are those of the authors and not necessarily those of the NHS or the Department of Health and Social Care.

## References

1. Bush, K., G. Jacoby, and A. Medeiros, A Functional Classification Scheme for b-Lactamases and Its Correlation with Molecular Structure. Antimicrob Agents Chemother, 1995. 39(6): p. 1211–1233.

2. Bush, K. and P.A. Bradford, beta-Lactams and beta-Lactamase Inhibitors: An Overview. Cold Spring Harb Perspect Med, 2016. 6(8).

3. Tehrani, K. and N.I. Martin, beta-lactam/beta-lactamase inhibitor combinations: an update. Medchemcomm, 2018. 9(9): p. 1439–1456.

4. Ohlin, B., et al., Piperacillin/Tazobactam Compared with Cefuroxime/ Metronidazole in the Treatment of Intra-abdominal Infections. Eur J Surg, 1999. 165: p. 875–884.

5. Viscoli, C., et al., Piperacillin-tazobactam monotherapy in high-risk febrile and neutropenic cancer patients. Clin Microbiol Infect, 2006. 12(3): p. 212–6.

6. Cooke, J., et al., Longitudinal trends and cross-sectional analysis of English national hospital antibacterial use over 5 years (2008-13): working towards hospital prescribing quality measures. J Antimicrob Chemother, 2015. 70(1): p. 279–85.

7. Bou-Antoun, S., et al., Descriptive epidemiology of Escherichia coli bacteraemia in England, April 2012 to March 2014. Euro Surveill, 2016. 21(35).

8. Lee, J., et al., The impact of the increased use of piperacillin/tazobactam on the selection of antibiotic resistance among invasive Escherichia coli and Klebsiella pneumoniae isolates. Int J Infect Dis, 2013. 17(8): p. e638–43.

9. Jamal, W.Y., M.J. Albert, and V.O. Rotimi, High Prevalence of New Delhi Metallo-beta-Lactamase-1 (NDM-1) Producers among Carbapenem-Resistant Enterobacteriaceae in Kuwait. PLoS One, 2016. 11(3): p. e0152638.

10. Papp-Wallace, K.M. and R.A. Bonomo, New beta-Lactamase Inhibitors in the Clinic. Infect Dis Clin North Am, 2016. 30(2): p. 441–464.

11. Wang, J., et al., Semi-rational screening of the inhibitors and β-lactam antibiotics against the New Delhi metallo-β-lactamase 1 (NDM-1) producing E. coli. RSC Advances, 2018. 8(11): p. 5936–5944.

12. Monogue, M.L., et al., Detection of Piperacillin-Tazobactam-Resistant/Pan-beta-Lactam-Susceptible Escherichia coli with Current Automated Susceptibility Test Systems. Infect Control Hosp Epidemiol, 2017. 38(3): p. 379–380.

13. Stainton, S.M., et al., Prevalence, patient characteristics and outcomes of a novel piperacillin/tazobactam-resistant, pan-beta-lactam-susceptible phenotype in Enterobacteriaceae: implications for selective reporting. Clin Microbiol Infect, 2017. 23(8): p. 581–582.

14. Baker, T.M., et al., Epidemiology of Bloodstream Infections Caused by Escherichia coli and Klebsiella pneumoniae That Are Piperacillin-Tazobactam-Nonsusceptible but Ceftriaxone-Susceptible. Open Forum Infect Dis, 2018. 5(12): p. ofy300.

15. Schechter LM, et al., Extensive Gene Amplification as a Mechanism for Piperacillin-Tazobactam Resistance in Escherichia coli. MBio, 2018. 9(2): p. e00583–18.

16. Lartigue, M.F., et al., Promoters P3, Pa/Pb, P4, and P5 upstream from bla(TEM) genes and their relationship to beta-lactam resistance. Antimicrob Agents Chemother, 2002. 46(12): p. 4035–7.

17. Zhou, K., et al., Piperacillin-Tazobactam (TZP) Resistance in Escherichia coli Due to Hyperproduction of TEM-1 beta-Lactamase Mediated by the Promoter Pa/Pb. Front Microbiol, 2019. 10: p. 833.

18. Hansen, K.H., et al., Resistance to piperacillin/tazobactam in Escherichia coli resulting from extensive IS26-associated gene amplification of bla_TEM-1_. J Antimicrob Chemother, 2019. 74(11): p. 3179–3183.

19. Nicoloff, H., et al., The high prevalence of antibiotic heteroresistance in pathogenic bacteria is mainly caused by gene amplification. Nat Microbiol, 2019. 4(3): p. 504–514.

20. Adler, M., et al., High fitness costs and instability of gene duplications reduce rates of evolution of new genes by duplication-divergence mechanisms. Mol Biol Evol, 2014. 31(6): p. 1526–35.

21. Harmer, C.J. and R.M. Hall, IS26-Mediated Precise Excision of the IS26-aphA1a Translocatable Unit. MBio, 2015. 6(6): p. e01866–15.

22. Harmer, C.J., R.A. Moran, and R.M. Hall, Movement of IS26-associated antibiotic resistance genes occurs via a translocatable unit that includes a single IS26 and preferentially inserts adjacent to another IS26. MBio, 2014. 5(5): p. e01801–14.

23. Harmer, C.J. and R.M. Hall, IS26-Mediated Formation of Transposons Carrying Antibiotic Resistance Genes. mSphere, 2016. 1(2).

24. Institute, C.a.L.S., M07 Methods for Dilution Antimicrobial Susceptibility Testing for Bacteria That Grow Aerobically, in 11th ed. CLSI Standard M07. 2018, Clinical and Laboratory Standards Institute: Wayne, PA.

25. Wick, R.R., et al., Unicycler: Resolving bacterial genome assemblies from short and long sequencing reads. PLoS Comput Biol, 2017. 13(6): p. e1005595.

26. Mikheenko, A., et al., Versatile genome assembly evaluation with QUAST-LG. Bioinformatics, 2018. 34(13): p. i142–i150.

27. Seemann, T., Prokka: rapid prokaryotic genome annotation. Bioinformatics, 2014. 30(14): p. 2068–9.

28. Wick, R.R., et al., Bandage: interactive visualization of de novo genome assemblies. Bioinformatics, 2015. 31(20): p. 3350–2.

29. Larsen, M.V., et al., Multilocus sequence typing of total-genome-sequenced bacteria. J Clin Microbiol, 2012. 50(4): p. 1355–61.

30. Joensen, K.G., et al., Rapid and Easy In Silico Serotyping of Escherichia coli Isolates by Use of Whole-Genome Sequencing Data. J Clin Microbiol, 2015. 53(8): p. 2410–26.

31. S, K., et al., Versatile and open software for comparing large genomes. Genome Biology, 2004. 5(2): p. R12.1–R12.9.

32. Lee, I., et al., OrthoANI: An improved algorithm and software for calculating average nucleotide identity. Int J Syst Evol Microbiol, 2016. 66(2): p. 1100–1103.

33. Zankari, E., et al., Identification of acquired antimicrobial resistance genes. J Antimicrob Chemother, 2012. 67(11): p. 2640–4.

34. Carattoli, A., et al., In silico detection and typing of plasmids using PlasmidFinder and plasmid multilocus sequence typing. Antimicrob Agents Chemother, 2014. 58(7): p. 3895–903.

35. Chung, C., S. Niemela, and R. Miller, One-step preparation of competent Escherichia coli: Transformation and storage of bacterial cells in the same solution. Proceedings of the National Academy of Sciences of the United States of America, 1989. 86: p. 2172–2175.

36. Thulin, M., BAT: an online tool for analysing growth curves. 2018: Retrieved from http://www.mansthulin.se/bat/.

37. Testing, T.E.C.o.A.S., Breakpoint tables for interpretation of MICs and zone diameters, in Version 10.0. 2020: http://www.eucast.org.

38. Tansirichaiya, S., M.A. Rahman, and A.P. Roberts, The Transposon Registry. Mob DNA, 2019. 10: p. 40.

39. Drawz, S. and R. Bonomo, Three Decades of β-Lactamase Inhibitors. Clinical Microbiology Reviews, 2010. 23(1): p. 160–201.

40. Harris, P.N.A., et al., Effect of Piperacillin-Tazobactam vs Meropenem on 30-Day Mortality for Patients With E coli or Klebsiella pneumoniae Bloodstream Infection and Ceftriaxone Resistance: A Randomized Clinical Trial. JAMA, 2018. 320(10): p. 984–994.

41. Sharara, S.L., et al., Is Piperacillin-Tazobactam Effective for the Treatment of Pyelonephritis Caused by ESBL-producing Organisms? Clin Infect Dis, 2019.

42. Weber, R.E., et al., IS26-Mediated Transfer of bla NDM-1 as the Main Route of Resistance Transmission During a Polyclonal, Multispecies Outbreak in a German Hospital. Front Microbiol, 2019. 10: p. 2817.

43. He, S., et al., Insertion Sequence IS26 Reorganizes Plasmids in Clinically Isolated Multidrug-Resistant Bacteria by Replicative Transposition. mBio, 2015. 6(3): p. e00762.

44. Bush, K. and P.A. Bradford, Interplay between beta-lactamases and new beta-lactamase inhibitors. Nat Rev Microbiol, 2019. 17(5): p. 295–306.

45. Vogwill, T., M. Kojadinovic, and R.C. MacLean, Epistasis between antibiotic resistance mutations and genetic background shape the fitness effect of resistance across species of Pseudomonas. Proc Biol Sci, 2016. 283(1830).

46. Hubbard, A.T.M., et al., Effect of Environment on the Evolutionary Trajectories and Growth Characteristics of Antibiotic-Resistant Escherichia coli Mutants. Front Microbiol, 2019. 10: p. 2001.

