## Supplementary Material for "Within-patient evolution to piperacillin/tazobactam resistance in a clinical isolate of *Escherichia coli* due to IS*26*-mediated amplification of *bla*_TEM-1B_"

**Supplementary Methods 1:**

*DNA extraction*

Genomic DNA for qPCR was extracted using the PureGene® Yeast/Bact kit B (Qiagen, Germany) following the manufacturer’s instructions for extraction of DNA from Gram-negative bacteria after normalising each culture to an optical density at 600nm (OD_600_) of 1 and eluted in molecular grade water.

Long fragment genomic DNA extraction for PCR amplification and both Illumina and ONT sequencing was performed using the Genomic DNA Buffer Set, following the manufacturer’s instructions for genomic DNA extraction using the Genomic-tip 100/G (both Qiagen, Germany) but eluted in molecular grade water.

Plasmid extraction was performed from 100 ml LB broth cultures with the Plasmid Midi Kit using Genomic-Tip 100 (both Qiagen, Germany) following the manufacturer’s instructions and eluted in molecular grade water.

Total DNA concentration was measured using a QuBit 4.0 Fluorometer (Invitrogen, US), with the dsDNA Broad Range kit.

*PCR*

All PCR reactions were performed using 0.02 U/µl Q5® High-Fidelity DNA Polymerase, 1x Q5® Reaction Buffer, 200 µM dNTPs and 0.5 µM of each primer (table S1) in a total volume of 50 µl.

16S rRNA PCR amplification was performed using the following protocol: denaturation at 98°C for 30 seconds, followed by 45 cycles of denaturation at 98°C for 10 seconds, annealing at 53°C for 30 seconds and elongation at 72°C for 30 seconds, followed by a single, final extension of 2 minutes at 72°C.

The detection of the RM/TU was performed using the following PCR protocol: denaturation at 98°C for 30 seconds, followed by 35 cycles of denaturation at 98°C for 10 seconds, annealing at 68°C (left junction of the composite transposon in the chromosome and TU) or 67°C (right junction of the composite transposon in the chromosome) for 30 seconds and elongation at 72°C for 17 seconds (left junction of the composite transposon in the chromosome), 25 seconds (right junction of the composite transposon in the chromosome) or 20 seconds (TU), followed by a single, final extension of 2 minutes at 72°C.

Detection of the formation of new junctions in the pHSG396:IS*26* due to the insertion of the TU using the following PCR protocol: denaturation at 98°C for 30 seconds, followed by 35 cycles of denaturation at 98°C for 10 seconds, annealing at 68°C (left junction insertion) or 67°C (right junction of the insertion) for 30 seconds and elongation at 72°C for 15 seconds (left junction) and 14 seconds (right junction), followed by a single, final extension of 2 minutes at 72°C.

The PCR products from the left side of the RM in the chromosome and the TU were PCR purified using the Monarch® PCR and DNA Clean-up Kit (New England Biolabs, USA) following the manufacturer’s protocol and eluted in molecular grade water. The PCR products from the right side of the RM in the chromosome and the left and right junction of the TU insertion into pHSG396:IS*26* were extracted from the gel using the Monarch® DNA Gel Extraction Kit (New England Biolabs, USA) following the manufacturer’s instructions and eluted in molecular grade water. All purified DNA extracts were then Sanger sequenced by GeneWiz (Takley, UK).

*IS*26 *plasmid construct*

IS*26* from 190963 (TZP-sensitive isolate) was amplified using 0.02 U/µl Q5® High-Fidelity DNA Polymerase, 1x Q5® Reaction Buffer, 200 µM dNTPs and 0.5 µM of the primers CMCap_F and CMCap_R (Table S1) in a total volume of 50 µl. Amplification was performed using the following protocol; Denaturation at 98°C for 30 seconds, followed by 35 cycles of denaturation at 98°C for 10 seconds, annealing at 69°C for 30 seconds and elongation at 72°C for 9 seconds, followed by a single, final extension step 72°C for 2 minutes.

The restriction sites for the restriction enzymes EcoRI and KpnI were added to the 5’ ends of the 820 bp double stranded amplicon by a second round of PCR, using the primers CMCapRE_F and CMCapRE_R (Table S1). Amplification was performed using the same protocol as the initial amplification of IS*26*, except with an annealing temperature of 72°C for 30 seconds.

The resulting PCR product and the recipient plasmid pHSG396 were digested with the restriction enzymes EcoRI and KpnI (both New England Biolabs, US). PCR clean-up was then performed using the Monarch PCR and DNA Clean-up Kit following the manufacturer’s instructions. The digested IS*26* PCR product and pHSG396 were ligated together using T4 ligase (New England Biolabs, US), at a 3:1 insert to vector mass ratio, incubated at room temperature for 10 minutes.

The ligated plasmid (5 µl) was transformed into NEB® 5-alpha competent *E. coli* (New England Biolabs, US) using the following protocol; the reaction was incubated on ice for 2 minutes, then heat shocked at 42°C for 30 seconds. Following a second incubation on ice for 2 minutes, 950 µL of SOC outgrowth medium was added and incubated at 37°C, 250 rpm for 1 hour. The transformed cells were plated out onto LB agar supplement with 35 µg/ml chloramphenicol and Isopropyl β-D-1-thiogalactopyranoside/X-gal (Fisher Scientific, USA) and incubated at 37°C for 18 hours and the plasmid was extracted following the protocol above. IS*26* in pHSG396 was Sanger sequenced by GeneWiz (Takely, UK) using commercially available M13 primers.

**Table S1:** Primer name and sequences used for PCR amplification and qPCR in this study

| Primer name | Primer Sequence |
| --- | --- |
| 16S-27F | AGAGTTTGATCCTGGCTCAG |
| 16S-1492R | TACCTTGTTACGACTT |
| RM2_F1 | TCTTCCCACTGCTGACGAAC |
| RM2_R1 | AGTGTGACGGAATCGTTGCT |
| RM2_F2 | CGCCCGAGATACAACATCCA |
| RM2_R2 | GTTGCTTTTTCTCGACGTGCT |
| CMCap_F | GGCACTGTTGCAAATAGTCGGT |
| CMCap_R | GGCACTGTTGCAAAGTTAGCGA |
| CMCapRE_F | AAAAAGAATTCGGCACTGTTGCAAATAGTCGGT |
| CMCapRE_R | AAAAAGGTACCGGCACTGTTGCAAAGTTAGCGA |
| CapCon_F1 | GGGGAAACGCCTGGTATCTT |
| CapCon_R1 | TGACGGAATCGTTGCTGTTG |
| CapCon_F2 | GCTGGGAATAGAACAGCCGA |
| CapCon_R2 | CGGCTCGTATGTTGTGTGGA |
| uidA_F | TCTGGCAACCGGGTGAAG |
| uidA R | TAGATATCACACTCTGTCTGGCT |
| TEM_F | GGAACCGGAGCTGAATGAAG |
| TEM_R | TCAGCAATAAACCAGCCAGC |
| OXA-1_F | TGCATCCACAAACGCTGAAA |
| OXA-1-_R | TGCGAAACCCAAACAACAGA |
| tetD_F | AGCAGAAACAAGAAAGCGCA |
| tetD_R | TTCTCACTCAGCCGTTTTGC |
| *aac*(3)*-lla_F* | GCATGCCTCACTTAAAGCGA |
| *aac*(3)*-lla_R* | ATTGATTCAGCAGGCCGAAC |
| *aac*(6’)*-lb-cr_F* | CCCAGTCGTACGTTGCTCTT |
| *aac*(6’)*-lb-cr_F* | CCTCGGGATCATTGAACAGC |

**Table S2:** Resistance profile of cefpodoxime (CPD), cefoxitin (FOX), piperacillin/tazobactam (TZP), meropenem (MEM), ciprofloxacin (CIP), cefotetan (CTT), amikacin (AMK), ertapenem (ETP), amoxicillin/clavulanic acid (AMC), chloramphenicol (CHL) and ampicillin (AMP) towards identified pair isolates with presumptive clonality assessed in clinic by disk diffusion method

| Patient Number | Isolate Number | CPD | FOX | TZP | MEM | CIP | CTT | AMK | ETP | AMC | CHL | AMP |
| --- | --- | --- | --- | --- | --- | --- | --- | --- | --- | --- | --- | --- |
| 66476 | 101737 | S | S | S | S | S | S | S | S | S | - | - |
|  | 102167 | S | S | **R** | S | **R** | S | S | S | I | - | - |
| 74418 | 153964 | S | S | S | S | S | S | S | S | **R** | S | **R** |
|  | 152025 | S | S | **R** | S | S | S | S | S | **R** | S | **R** |
| 90508 | 190693 | S | S | S | S | **R** | **R** | S | S | **R** | S | **R** |
|  | 169757 | S | S | **R** | S | **R** | **R** | S | S | **R** | S | **R** |

**Figure S1:** Restriction fragment length polymorphism of **A**) 16S rRNA PCR amplicon of **1**) 151985, **2**) 153006, **3**) 101737, **4**) 102167, **5**) 153964, **6**) 152025, **7**) 155512, **8**) 154810, **9**) 190693 and **10**) 169757 digested with AlwNI, PpuHI and MslI. RFLP of **B**) long fragment genomic DNA extraction from **11**) 153964, **12**) 152025, **13**) 190693 and **14**) 169757 digested with SpeI and MslI

Figure S1:

**A**

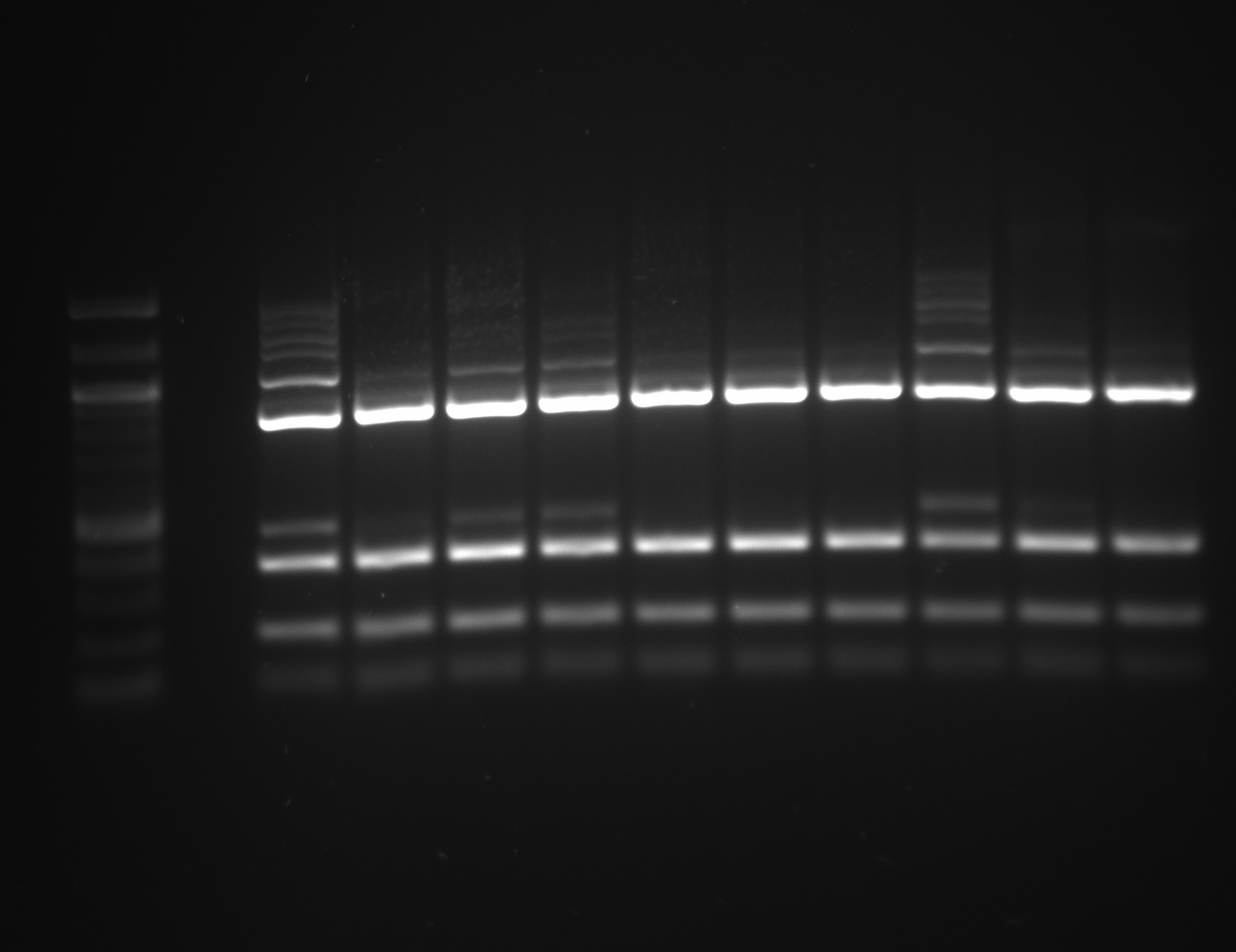

**1	2	3	4	5	6	7	8	9	10**

1517

1200

1000

900

800

700

600

500/517

400

300

200

100

**B**

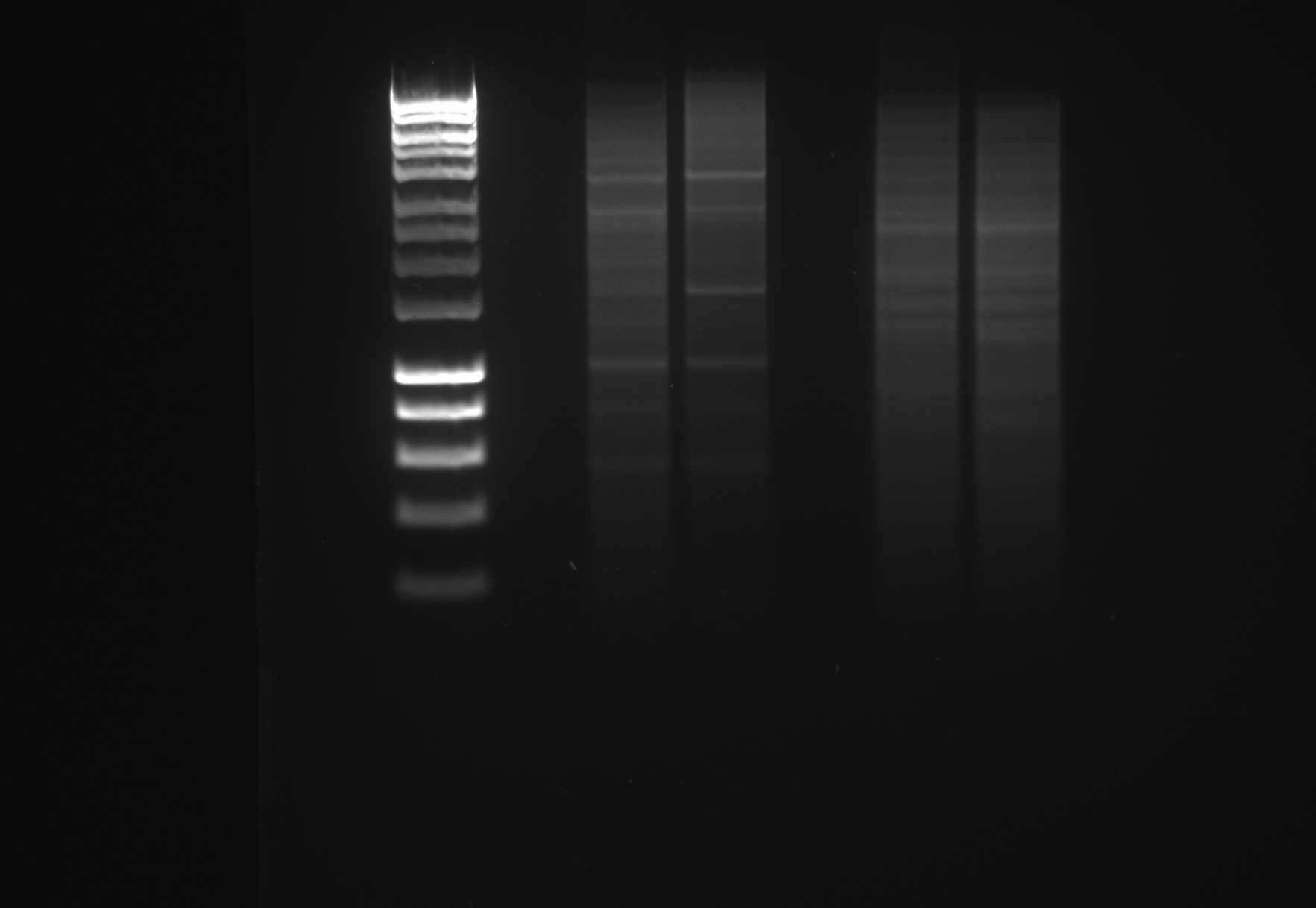

**11	12	13	14**

10037

8000

6000

5000

4000

3000

2500

2000

15001517

1000

800

600

400

200

**Figure S2:** Gel electrophoresis of the PCR amplicons of the **1**) left junction of the chromosomally located IS*26* and resistance module, **2**) right junction of the chromosomally located IS*26* and resistance module and **3**) translocatable unit from **A**) TZP-sensitive isolate and **B**) TZP-resistant isolate

Figure S2:

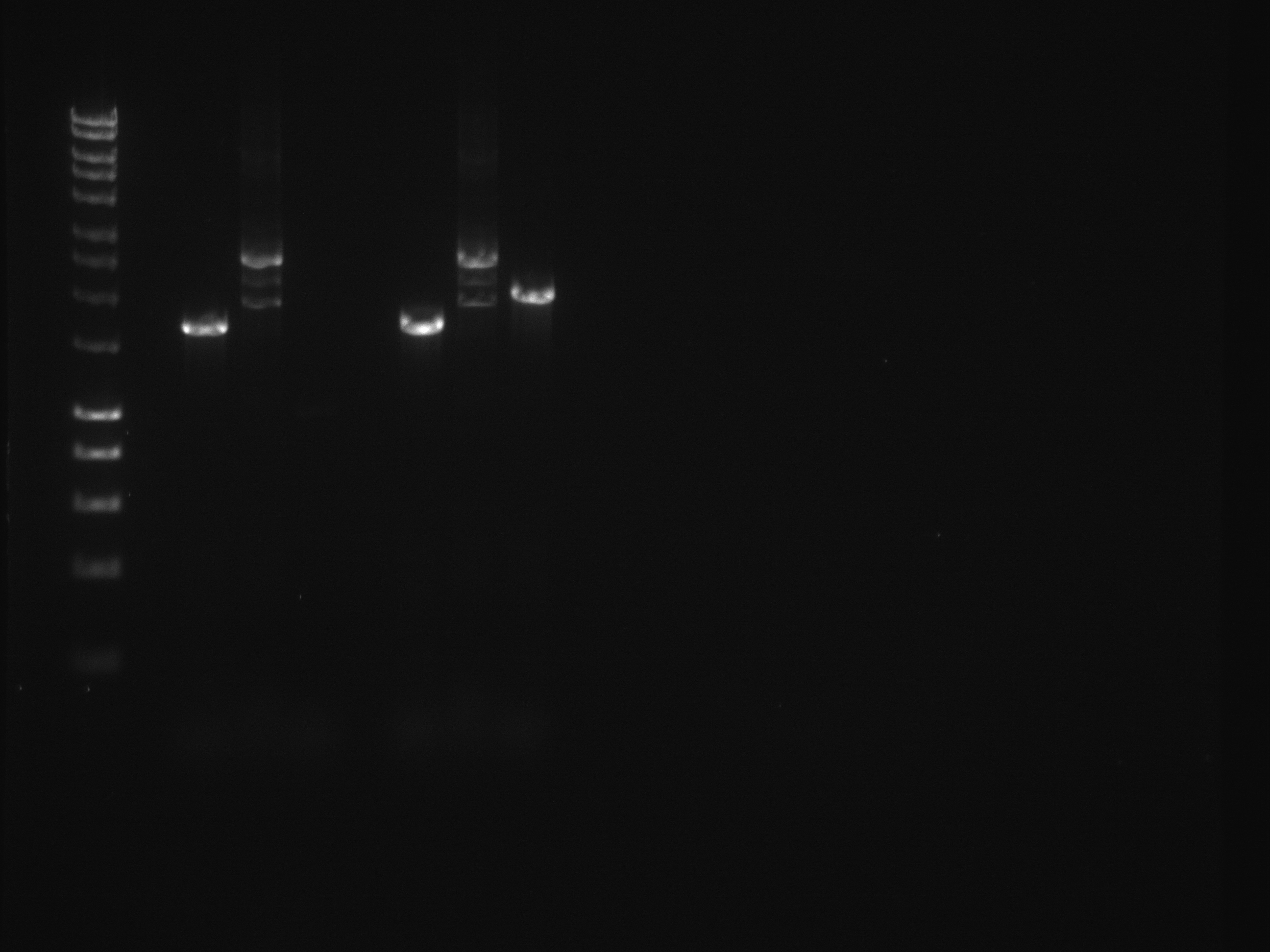

**1	2	3**

10037

8000

6000

5000

4000

3000

2500

2000

1500/1517

1000

800

600

400

200

**1	2	3**

**A**

**B**

**Figure S3:** **A**) Gel electrophoresis of **1**) undigested pHSG396:IS*26*, **2**) digested pHSG396:IS*26*, **3**) undigested pHSG396:IS26 after growth in the presence of 8/4 µg/ml TZP and transformation in NEB® 5-alpha competent *E. coli* and **4**) digested pHSG396:IS26 after growth in the presence of 8/4 µg/ml TZP and transformation in NEB® 5-alpha competent *E. coli* and **B**) gel electrophoresis of PCR amplicons from the **5**) left and **6**) right junctions of pHSG396:IS*26* following insertion of the TU

Figure S3:

**A**)

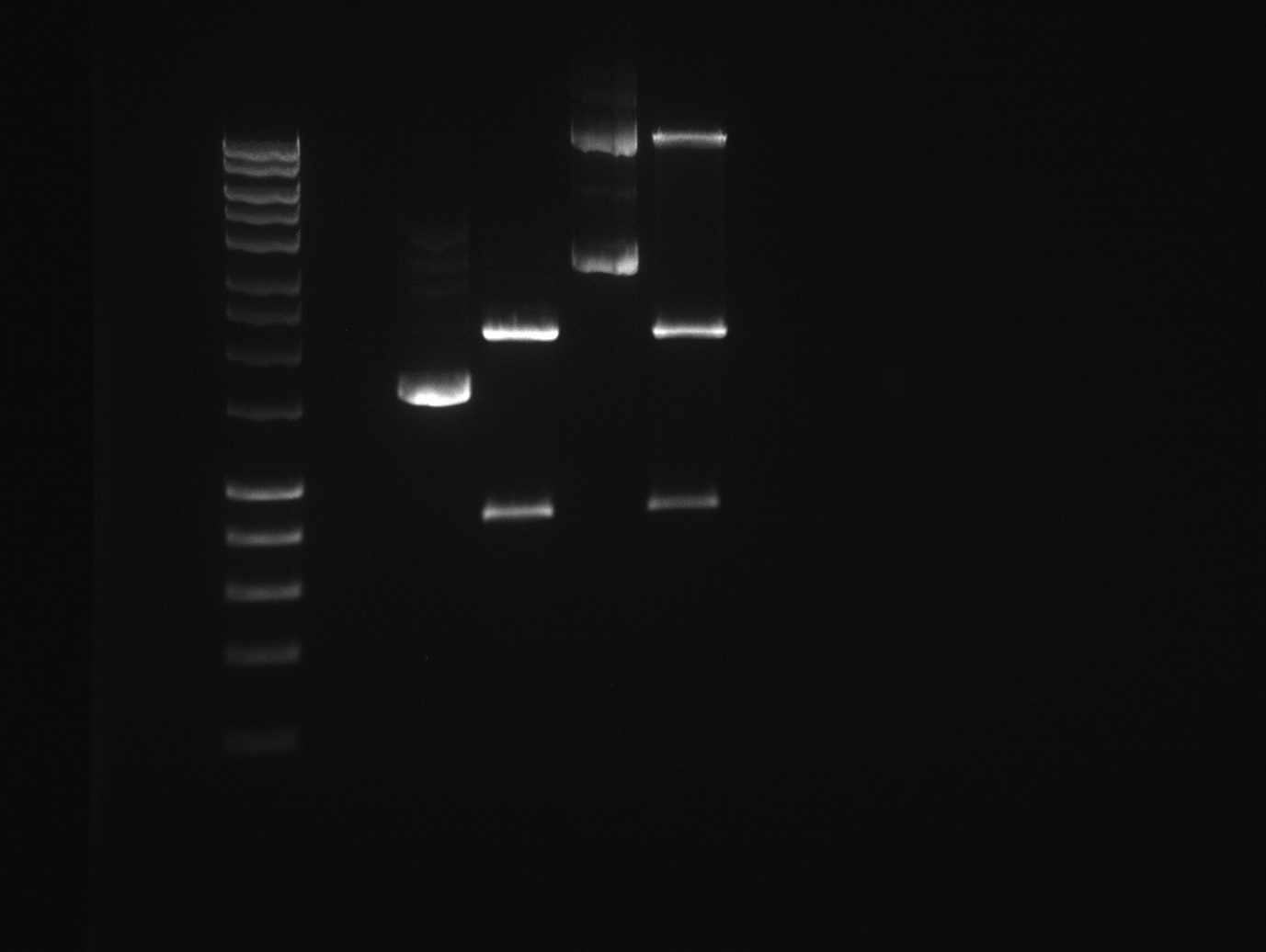

10037

8000

6000

5000

4000

3000

2500

2000

1500/1517

1000

800

600

400

200

1	2	3	4

**B**)

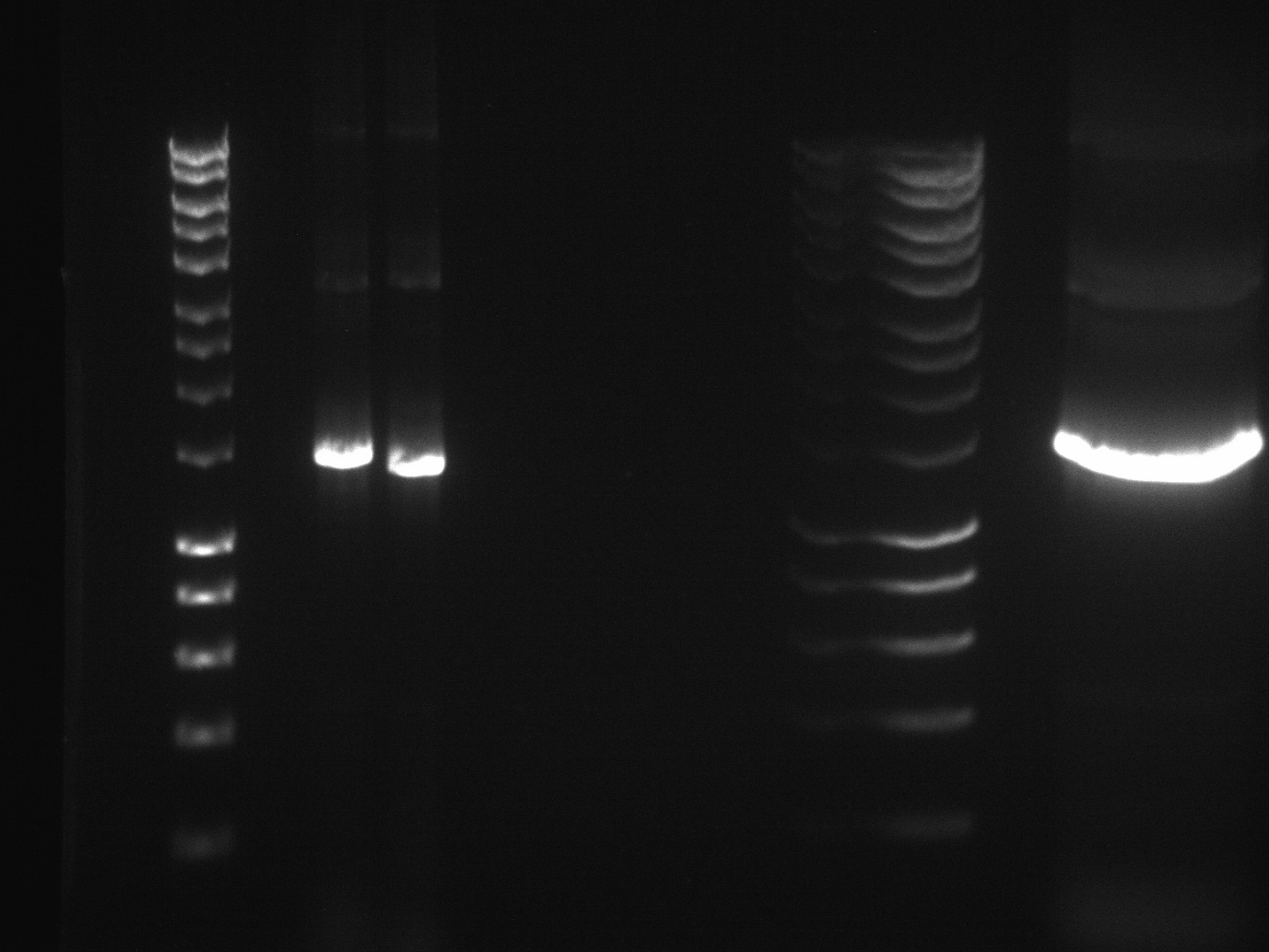

**5	6**

10037

8000

6000

5000

4000

3000

2500

2000

1500/1517

1000

800

600

400

200

**Figure S4:** Visualisation of the hybrid assembly of long and short read sequencing reads by Unicycler of the **A**) TZP-susceptible isolate and **B**) and TZP-resistant isolate

Figure S4:

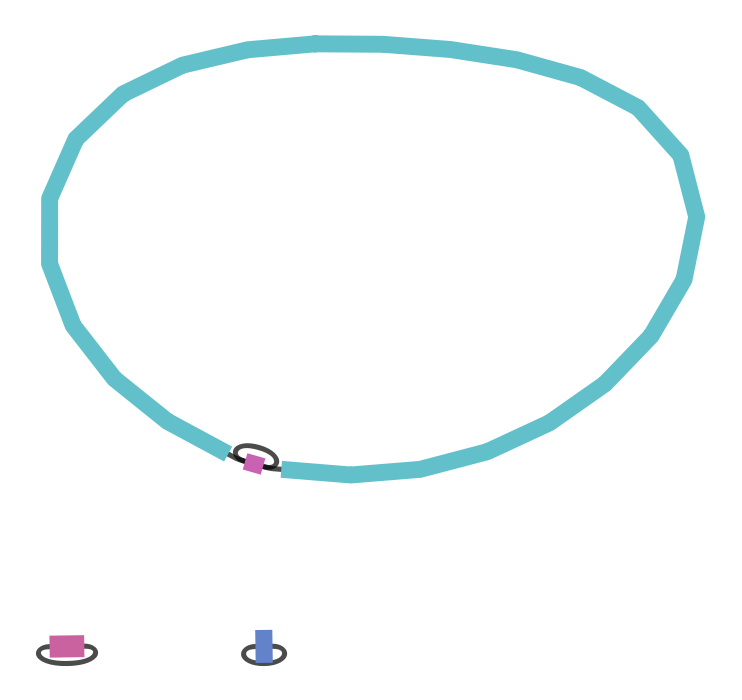

2.59x

106637bp

8.51x

10899bp

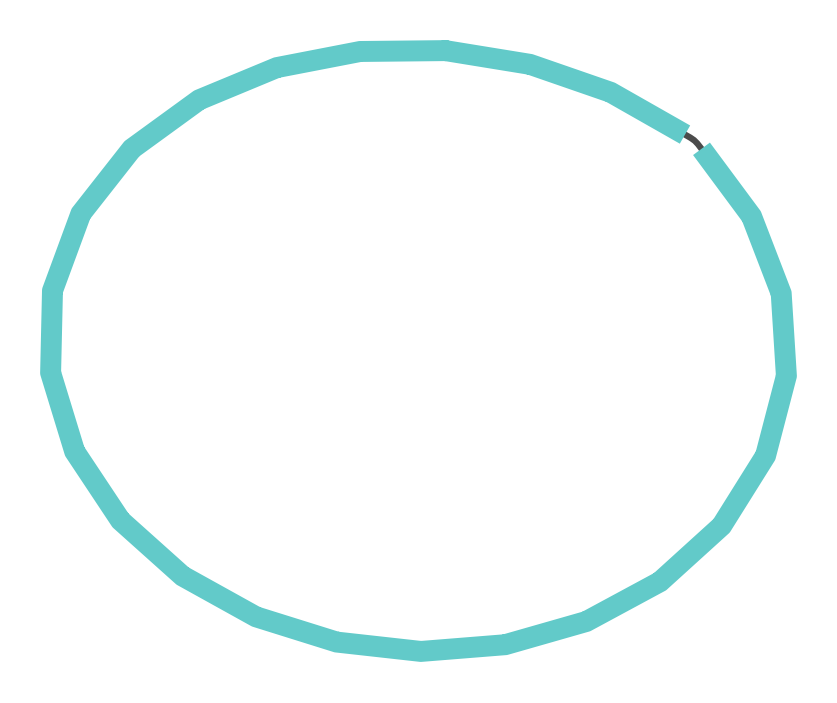

1x

5141277bp

1x

5151950bp

**A**

**B**
